# Pan-repository analysis reveals a drug-activating function of microbial bile acid conjugation

**DOI:** 10.64898/2026.03.03.709330

**Authors:** Vincent Charron-Lamoureux, Patricia Kelly, Simone Zuffa, Abubaker Patan, Marta Sala-Climent, Corinn Walker, Haoqi Nina Zhao, Shipei Xing, Harsha Gouda, Julius Agongo, Erin R. Reilly, Lina Sallam, Selene F.H. Shore, Subhomitra Ghoshal, Anika K. Harpavat, Marudhu Pandiyan Murugesan, Swara Yadav, James Versalovic, Vlad Orlovsky, Yasin El Abiead, Kine Eide Kvitne, Janet G. Markle, Grant J. Norton, Gregory T. Walker, Michael H. Lee, Zhewen Hu, Marvic Carrillo Terrazas, David M. Zong, Amir Zarrinpar, Manuela Raffatellu, Anthony Martin, Loryn Chin, Suzanne Devkota, Antonio Gonzalez, Gail Ackermann, Lucas Patel, Yuhan Weng, Rob Knight, Richard K. Russell, Richard Hansen, Vaios Svolos, Konstantinos Gkikas, Nicholas J.W. Rattray, Dionicio Siegel, Karsten Zengler, Maribeth R. Nicholson, Hiutung Chu, Andrew D. Patterson, Monica Guma, Richard Kellermayer, Melinda A. Engevik, Thomas D. Horvath, Konstantinos Gerasimidis, Pieter C. Dorrestein

**Author notes:** Overall project, data science of the bile acids and discovery (Vincent Charron-Lamoureux, Pieter C. Dorrestein). For the iPENS study (Konstantinos Gerasimidis). For the quantitation data (Thomas D. Horvath). For the mouse interventional study (Melinda A. Engevik). For the rheumatoid arthritis study (Monica Guma). Author contributions VCL and PCD conceptualized the project. VCL, PK, SZ, AG, LP, YW data analysis. KEK, JGM, MSC, HNZ, SX, HG, GTW, MHL, YEA, RKR, RH, VS, KG, NJWR, MG, DZ, AZ, and MR provided data or assisted with data analysis. VCL, GN, and CW performed culturing experiments under KZ supervision. VCL and JA acquired the LC-MS/MS data. AP, ZH, did the synthesis part of the project supervised by DS. TDH, AKH, MPM, SY, and VO did the LC-MS/MS quantitation and JV provided guidance and funding. MAE, SFHS, and SG generated the in vivo mouse experiments. RM and MRN collected samples from patients. ERR and LS did the PPAR-γ assay supervised by ADP. VCL and PCD wrote the manuscript and all authors reviewed, edited, and approved the manuscript. RK, MG, MRN, RKR, RH provided review and editing from the clinical perspective. Disclosures PCD is an advisor and holds equity in Cybele, BileOmix, Sirenas and a scientific co-founder, advisor, holds equity and/or received income from Ometa, Enveda, and Arome with prior approval by UC San Diego. PCD. also consulted for DSM animal health in 2023. TDH is a member of the Editorial Advisory Board and is contracted as an Associate Academic Editor for a Cell Press journal called STAR Protocols. TDH is also a participant in the SCIEX Global Thought Leaders in Mass Spectrometry Program. VO is a founder and CEO of Helix Chromatography. KZ is a scientific co-founder, advisor, holds equity and/or received income from Isolation Bio, Native Microbials, and Guilden with prior approval by UC San Diego. MG has research agreements with AbbVie and Janssen Pharmaceutica. KGk PhD studentship was funded in part by Nestle Health Science. In the last 3 years, Richard Russell received consultation fees, research grants, royalties, or honorarium from Johnson & Johnson, Pfizer, Ferring, Celltrion, Lilly, AlfaSigma & Pharmacosmos. VS received research/travel grants or consulting fees from NATPROD IKE, Mendes S.A., MDPI, SYN Innovation Laboratories S.A., Chronicles Health and Access Nutrients Inc. AZ is a co-founder and equity holder in Endure Biotherapeutics. JV is a scientific advisor for Seed Health. RK is a scientific advisory board member, and consultant for BiomeSense, Inc., has equity and receives income. RK is a scientific advisory board member and has equity in GenCirq. RK has equity in and acts as a consultant for Cybele. RK is a Vice President and board member of Microbiota Vault, Inc. He is a board member of N=1 IBS advisory board and receives income. RK is a Senior Visiting Fellow of HKUST Jockey Club Institute for Advanced Study. The terms of these arrangements have been reviewed and approved by the University of California, San Diego in accordance with its conflict of interest policies. PCD and VCL have filed a patent application on this material.

## Abstract

Microbially modified bile acids shape host physiology by regulating nutrient absorption, glucose homeostasis, circadian rhythms and thermoregulation. Here we identify a previously unrecognized drug-activating function of microbial bile acid conjugation. By systematically mining human LC–MS/MS datasets across public repositories and linking uncharacterized bile acid spectra to health-associated metadata, we discovered conjugates of the >75-year-old anti-inflammatory drug 5-aminosalicylic acid (5-ASA) with primary and secondary bile acids, including cholic, deoxycholic and lithocholic acids. These bile acid–drug conjugates were detected specifically in individuals treated with 5-ASA or its prodrugs. Multiple gut bacteria, including members of the Bacteroidota and Bacillota, generated cholyl–5-ASA in vitro, and bile salt hydrolase–associated transaminase activity was required for conjugate formation. In a mouse model of colitis, cholyl–5-ASA was associated with reduced intestinal inflammatory pathology and showed markedly enhanced activation of PPAR-γ in cell-based reporter assays compared with 5-ASA alone. Consistent with this activity, cholyl–5-ASA elicited selective immunophenotypic changes in CD4⁺ T cells in vitro, including increased Foxp3+ regulatory T cells. Together with prior evidence that 5-ASA efficacy depends on the microbiome, these findings support a model in which microbial bile acid conjugation represents a key activation step for 5-ASA therapy. More broadly, this work demonstrates how pan-repository metabolomics can uncover previously unrecognized microbiome-dependent chemical functions with direct therapeutic relevance.

## Main

Bile acids play critical roles in human physiology and have been implicated in diverse health outcomes, including cancer^1,2^, Alzheimer’s disease^3^, atherosclerosis^4^, metabolic dysfunction-associated steatohepatitis^5^, obesity^6^, diabetes^7^, sleep disruption^8^, inflammatory bowel disease (IBD)^9^, and chronic pruritis^10^. Yet our understanding of bile acid biology has been largely shaped by a narrow subset of ∼20 commercially available compounds used for clinical diagnostics. The advent of modern mass spectrometry-based metabolomics, coupled with scalable data science infrastructure, is now revealing a diversity of bile acids that was unimaginable just a few years ago, including microbial modifications of the carboxylate (amidates) that were previously overlooked^5,7,11–20^.

Despite being among the first biomolecules structurally characterized - with the first bile acid structure reported in 1848^21^ and taurine and glycine conjugates described in the 1930s - our understanding of the structural diversity of bile acids has experienced unprecedented growth in the last few years. Modifications such as ornithine, arginine, and other amino acid and dipeptide conjugates were noted sporadically in the 1960–1980s, but were largely ignored until our 2020 study revealed that gut microbes produce additional amino acid amidates, including Phe, Tyr, Leu, and others, whose levels are increased in IBD and associated with immune regulatory functions^11^. While microbial bile acid amidates are now recognized as modulators of early life physiology^7,13^, aging^14^, and disease, including IBD^11,15^, diabetes^7,16,17^, and colorectal cancer^2^, the integration of comprehensive bile acid network mapping across medical specialties is in its infancy. Leveraging single repository-scale liquid chromatography-tandem mass spectrometry (LC-MS/MS) data analysis (GNPS-MassIVE), we identified 21,549 MS/MS spectra, spanning multiple ion forms, multimers, and in-source fragments, representing thousands of yet-uncharacterized bile acids^12^. To link bile acids to health conditions, we performed large-scale MS/MS searches with the bile acid MS/MS library across GNPS/MassIVE^18^, MetaboLights^22^, and Metabolomics Workbench^23^ using FASST^24^, a rapid implementation of MASST^25^, and integrated the results with disease annotations through harmonized Pan-ReDU metadata^26,27^. This approach reveals an expansive hypothetical bile acid-health condition landscape and provides a framework for connecting molecular diversity to human health at a scale that is still largely underappreciated in bile acid biology. This analysis led to the discovery of a cholic acid conjugate of 5-aminosalicylic acid (5-ASA), an anti-inflammatory drug bile acid conjugate, observed in humans and mice that we show is active in suppressing the induction of colitis in mice.

## Results

### Analysis of candidate bile acid diversity across human health conditions

We linked 21,549 previously uncovered bile acid MS/MS spectra to LC–MS/MS datasets with harmonized disease-ontology metadata. To achieve this, we searched the bile acid MS/MS library against ∼1.3 billion pan-repository indexed MS/MS spectra (as of February 2025) using FASST, spanning datasets deposited in metabolomics repositories. The analysis included LC-MS/MS datasets from across biospecimen types (e.g., stool, blood, urine, and tissues) with disease and health status information using pan-ReDU metadata^26^. Pan-ReDU links LC-MS/MS data to sample information encompassing 93 body sites/biofluids and 25 health conditions. In total, we obtained pan-repository MS/MS spectral matches for 241,752 human LC–MS/MS files available at the time of this analysis, of which 137,776 contained health-related metadata. Within this repository-scale analysis, 76 spectral matches unique to diabetes and obesity datasets, including N-acetylglucosamine–derived conjugates discovered in 1989^28^, while 48 spectra were associated with rheumatoid arthritis (RA) and IBD (Fig. 1). Notably, only six MS/MS spectra were observed across all major IBD categories - whether labeled simply as “IBD” or specifically as ulcerative colitis (UC) or Crohn’s disease (CD) - indicating that these results spanned multiple independent datasets and provide a first clue that this signal may be IBD specific (Fig. 1). These six spectra yielded a total of 634 matches across the pan-repository data. The putative modifications corresponding to these shared MS/MS spectra included mass differences - relative to unconjugated bile acids - of Δ*m/z* 135.0319 (whose MS/MS is consistent with a trihydroxylated bile acid core^12^), Δ*m/z* 117.0211 (linked to di-and tri-hydroxylated bile acids), Δ*m/z* −18.99 (and bile acid MS/MS signatures that match a monohydroxylated bile acid), and Δ*m/z* 390.2770 (trihydroxylated bile acid). The mass difference between the Δ*m/z* 135.0319 and Δ*m/z* 117.0211 species corresponded to a water loss, suggesting that the Δ*m/z* 117.0211 and the Δ*m/z* 135.0319 containing species are related by a water loss that could be an in-source fragment. We therefore focused our attention on the Δ*m/z* 135.0319 species.

**Fig. 1.**
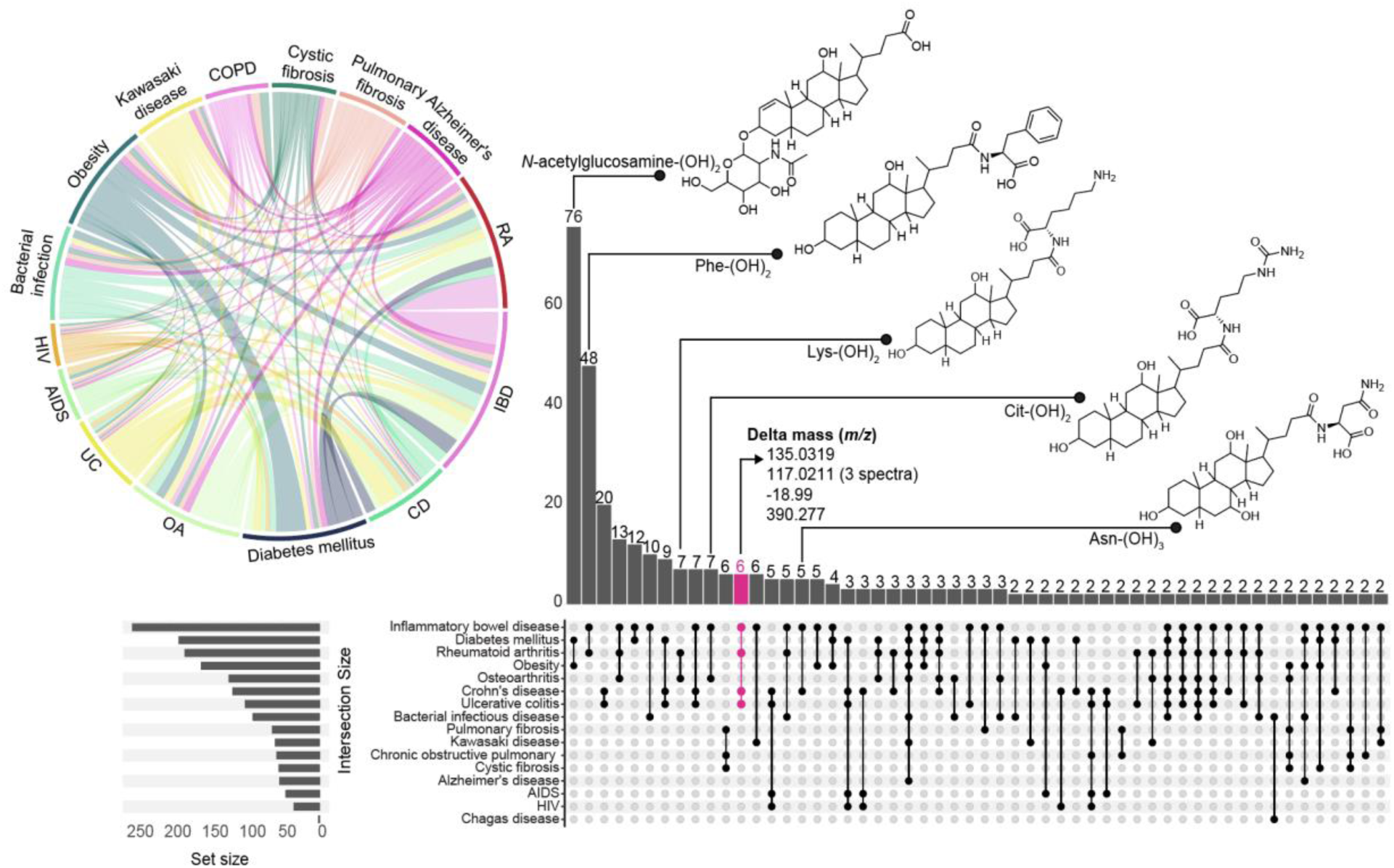
Health associations derived from 23,656 human-only MS/MS spectra matching bile acid fragmentation patterns across metabolomics repositories GNPS/MassIVE, MetaboLights, and Metabolomics Workbench that have harmonized metadata. UpSet plot illustrating the number of bile acid MS/MS shared across multiple health conditions with data indexed as of February 2025. Each vertical column represents the number of shared MS/MS spectra of candidate bile acids, which is connected to the disease information represented by the column on the left, called set size. Chord diagram shows the distribution of bile acids among diseases. Structures are representative bile acids for which MS/MS of standards are matched. CD, Crohn’s disease; IBD, inflammatory bowel disease; RA, rheumatoid arthritis; COPD, chronic obstructive pulmonary disease; HIV, human-immunodeficiency virus; AIDS, acquired immunodeficiency syndrome; UC, ulcerative colitis; OA, osteoarthritis. The structures shown are example annotations.

### Discovery and confirmation of the existence of 5-ASA drug–bile acid conjugates

As the Δ*m/z* 135.0319 was an uncharacterized bile acid modification, we set out to understand its complete structure. Outside of IBD, we also found the same uncharacterized bile acids in data from people with RA. This was unexpected given that one is an intestinal immune-mediated inflammatory disease and the latter is an extraintestinal autoimmune disorder. The stool extracts from the RA study were available in our lab and this dataset, which was composed of samples collected from 20 individuals designed to explore an anti-inflammatory diet, had been previously published^29^. These data and the reanalysis of the sample extracts were instrumental to our understanding of what the Δ*m/z* 135.0319 bile acid modification was.

Molecular networking, a method that aligns MS/MS spectra based on spectral similarity, clustered the Δ*m/z* 135.0319 features with known bile acids such as phenylalanine-dihydroxylated bile amidates, glycine bile amidates, and several other bile acids with unknown structures supporting shared core structural similarities with known bile acid amidates (Fig. 2a). Incorporating patient metadata information, including their medication history (e.g., methotrexate, sulfasalazine, hydroxychloroquine, and leflunomide), we found three metabolic features (*m/z* 544.3269, *m/z* 528.3342, *m/z* 512.3373), each with a Δ*m/z* 135.0319 (molecular formula C_7_H_5_NO_2_) compared to the unmodified bile acids. Strikingly, these features were detected exclusively in two individuals (patients 28 and 4) (Fig. 2b). Cross-referencing their medical records showed that both were receiving sulfasalazine, a disease-modifying antirheumatic drug (DMARD) prescribed in RA. In this prodrug, the 5-ASA component is considered inactive, while the sulfapyridine component, released during metabolism, is regarded as the active therapeutic portion^30–34^. Thus, in this cohort, the two people that took sulfasalazine also had a potential bile acid - drug conjugate in the stool, while 18/18 people who did not take the drug also did not have these bile acid conjugates. The probability of this finding being a random connection is unlikely with an odds ratio of 185 (Haldane–Anscombe corrected), 95% CI: 2.9 - 11,740, Fisher’s exact *p*-value: 0.0053. Sulfasalazine is a prodrug relying on a microbial azoreductase to cleave the azo bound releasing sulfapyridine and 5-aminosalicylic acid (5-ASA)^35^, which has a Δ*m/z* 135.0319 (after dehydration from the condensation reaction based conjugation to bile acids). Therefore, we hypothesized that 5-ASA (molecular formula: C_7_H_7_NO_3_) was the molecule that we detected conjugated to cholic acid (*m/z* 544.3269), to deoxycholic acid (*m/z* 528.3342), and to lithocholic acid (*m/z* 512.3373) (Fig. 2c).

**Fig. 2.**
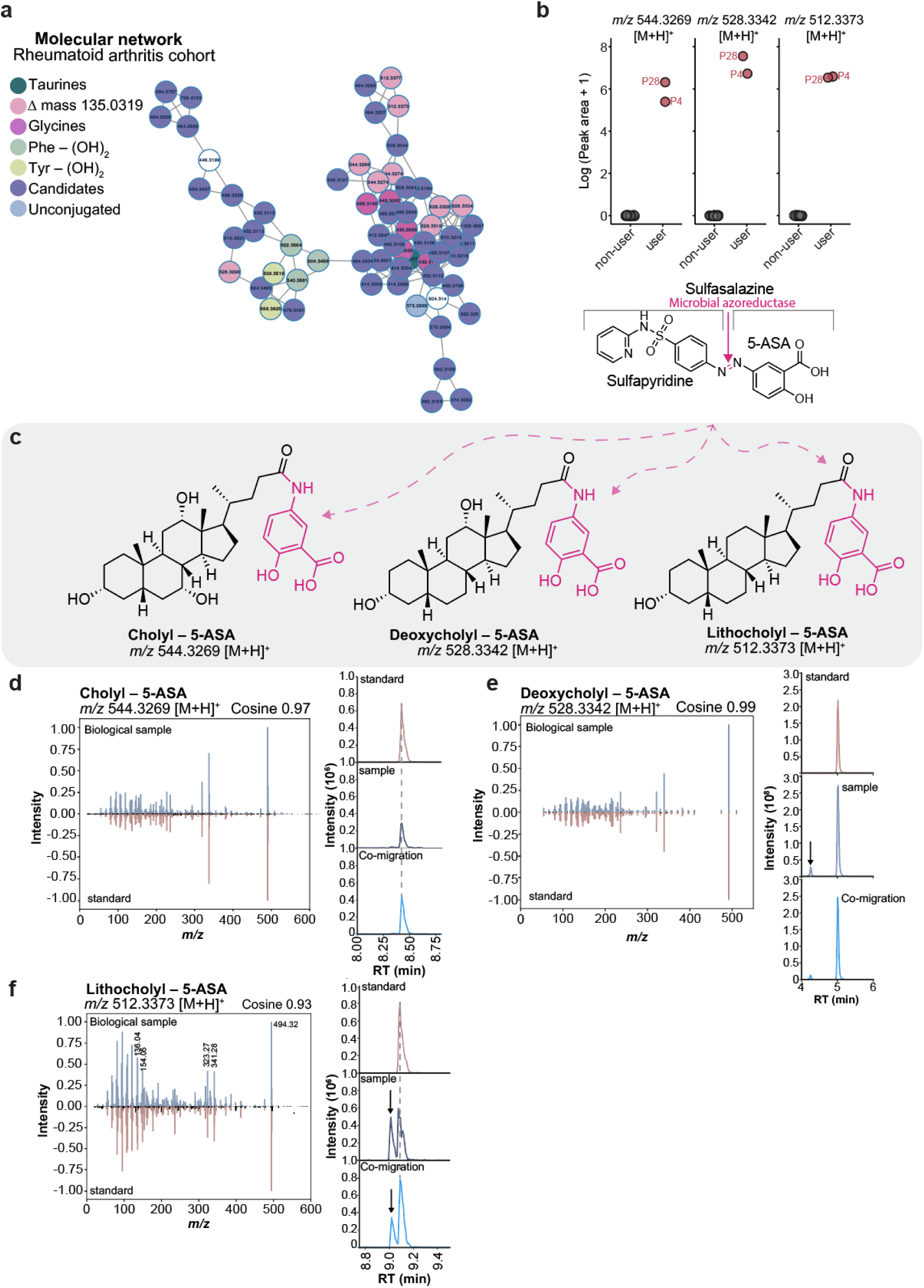
Discovery of 5-aminosalicylic acid-bile acid conjugates in a rheumatoid arthritis cohort. a, Molecular network of bile acids found in the stool of individuals with rheumatoid arthritis, highlighting the clustering of the putative bile acids with Δ*m/z* 135.0319 (pale pink) with taurine conjugates (teal), glycine conjugates (violet), microbial conjugated bile acids with phenylalanine (Phe-(OH)2, sage), microbial conjugate bile acids with tyrosine (Tyr-(OH)2, lime), other unknown bile acids (dark blue), and unconjugated bile acids (blue). **b,** Peak areas of features matching *m/z* 544.3269, *m/z* 528.3342, and *m/z* 512.3373, (with a Δ*m/z* 135.0319) are exclusively observed in two patients (Patient 4 and Patient 28), both reporting the use of the prodrug sulfasalazine. Sulfasalazine is activated via microbial azoreductase releasing sulfapyridine and 5-ASA. **c,** Proposed structures of the drug 5-ASA conjugated to cholic acid, deoxycholic acid, and lithocholic acid. **d-f,** Mirror plot showing MS/MS spectral similarity between the synthetic standard and the biological sample and extracted ion chromatogram from the standard, biological sample, and spiked-in of cholyl–5-ASA, deoxycholyl–5-ASA, and lithocholyl–5-ASA. The arrows denote related isomers of lithocholic acid and deoxycholic acid.

To validate the identities of these molecules, we synthesized pure synthetic standards of the cholic acid, deoxycholic acid, and lithocholic acid conjugates of 5-ASA and compared their retention time and MS/MS spectra with the biological samples. All compounds matched both the retention time and spectral similarities with the synthetic standards, confirming their presence in stool samples (Fig. 2d-f). It should be noted that we also detected other monohydroxylated and dihydroxylated 5-ASA conjugates present in the RA stool sample extracts that have different retention times compared to the standards we synthesized. While these features are consistent with bile acid–5-ASA conjugates, their exact identities remain to be elucidated.

### Cholyl–5-ASA is observed in public data across six independent IBD cohorts

Sulfasalazine is the only 5-ASA-containing prodrug approved for use in RA, and its use is infrequent in this indication compared to IBD. In addition, 5-ASA as mesalazine and related prodrugs, such as balsalazide and olsalazine, are prescribed for IBD, particularly UC. All prodrugs release 5-ASA upon activation. 5-ASA is used as both an induction and a maintenance therapy in patients with mild colitis. We thus hypothesized that 5-ASA also undergoes these same bile acid conjugations in patients with IBD, if 5-ASA is indeed modified with bile acids and would allow for additional confirmation that this bile acid conjugate exists.

To investigate the prevalence of cholyl–5-ASA across IBD data for which 5-ASA and its prodrugs (e.g., sulfasalazine, balsalazide, olsalazine) are prescribed, we leveraged the FASST search engine to find public metabolomics datasets containing MS/MS spectral matches to cholyl–5-ASA. We then prioritized datasets with curated metadata, including patient medication histories, identifying three adult IBD cohorts and one paediatric IBD cohort for downstream analysis. It should be noted that none of the studies were designed to specifically evaluate the efficacy of 5-ASA medications. Therefore, the reanalysis is only leveraged to assess if known 5-ASA medications use are linked to the formation of the bile acid conjugations in humans.

The first cohort included 197 stool samples (21 healthy, 59 UC, 117 CD; Fig. 3a). 5-ASA exposure was categorized as current, prior, non-users, and unknown^36^. Consistent with findings across all cohorts analyzed within this study, whose MS/MS matches to cholyl–5-ASA was not detected in any healthy individuals. Among 31 subjects with UC reporting current 5-ASA use, 5-ASA was detectable in 23 individuals. Cholyl–5-ASA was observed in 17 of these 23 individuals (74%) and in 6 of the prior users (26%) consistent with reduced conjugate detection after discontinuation (Fig. 3a). We could not do a correlation between last dose and time of sample collection as this information was not available. Among 117 subjects with CD, 5-ASA was detected in 36 individuals, and cholyl–5-ASA was observed in 13 of them (36%; Supplementary Fig. S1a). Medication use metadata were not captured for the CD patients, however, 5-ASA exposure could be directly evaluated from metabolomics data using a recently developed drug library^37,38^.

**Figure 3.**
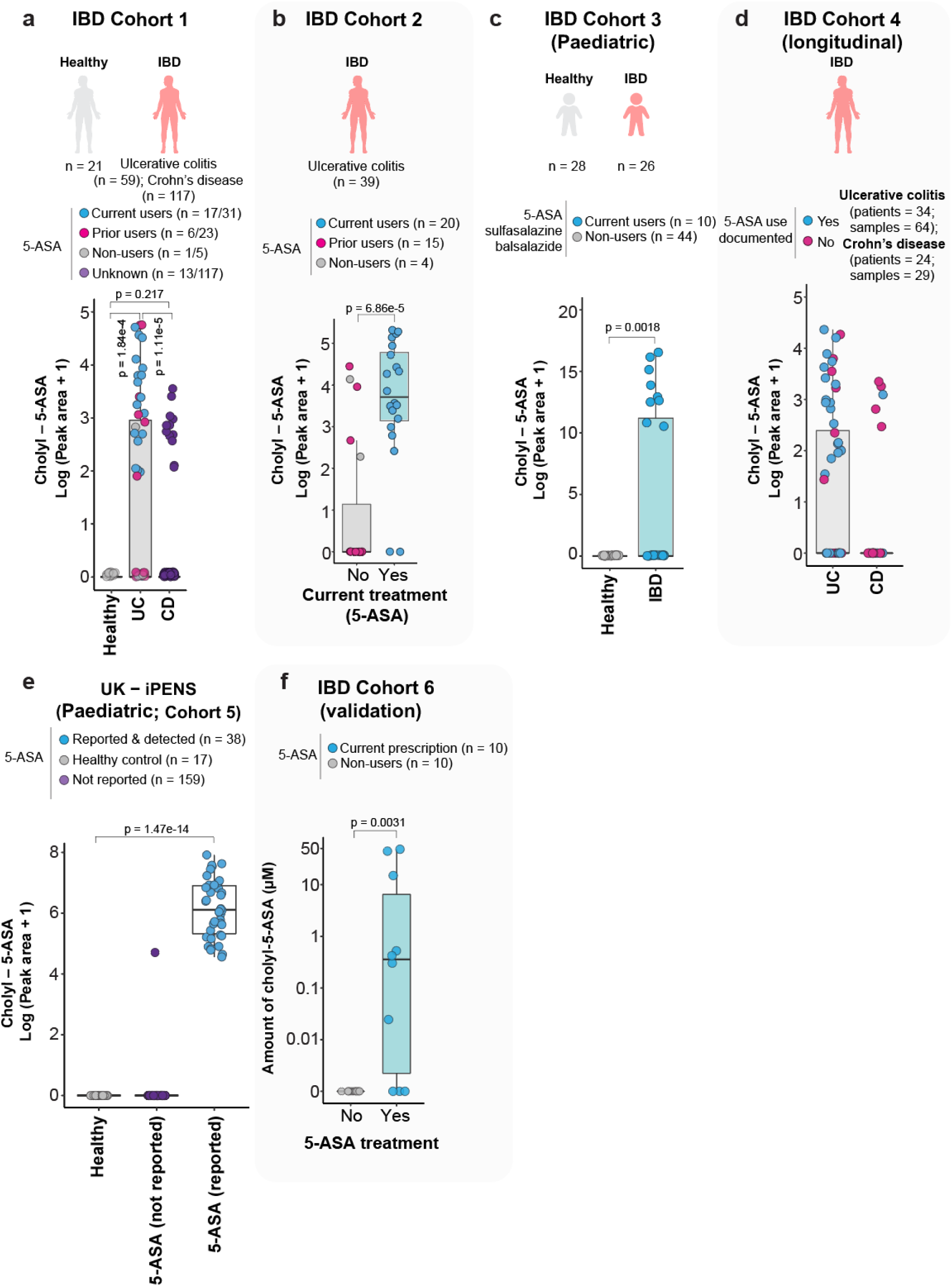
Cholyl–5-ASA is observed across six different IBD cohorts. a, Extracted peak area of the mass spectrometry features with the MS/MS of cholyl-5-ASA in healthy volunteers (n = 21), people with ulcerative colitis (n = 59), and Crohn’s disease (n = 117). 5-ASA exposure types were color-coded; current users (blue), prior users (pink), non-users (gray), and unknown (purple). Numbers in parentheses represent the number of samples in which cholyl–5-ASA is detected over the total number of samples available per IBD disease subtypes. **b,** Extracted peak area of cholyl–5-ASA in current treatment of 5-ASA of 40 ulcerative colitis patients. 5-ASA exposure types were color-coded; current users (blue), prior users (pink), non-users (gray). **c,** Extracted peak area of cholyl–5-ASA in healthy (n = 28) and IBD (n = 26) of a paediatric cohort of IBD. 5-ASA exposure types were color-coded; current users (blue) and non-users (gray). **d,** Extracted peak area of cholyl–5-ASA from a longitudinal study of individuals with ulcerative colitis (n = 34; samples 64) and Crohn’s disease (n = 24; samples 29) filtered for detectable 5-ASA from a IBD surgery outcome survey. Points are color-coded by 5-ASA exposure; documented 5-ASA use (blue) and not documented use (pink). **e**, Detection of cholyl–5-ASA in the untargeted data from the UK iPENS study. The n represents samples. **f**, Concentration (µM) of cholyl–5-ASA in 10 individuals without 5-ASA prescriptions and 10 individuals undergoing 5-ASA treatment. Significance tested via Fisher’s exact test.

The second cohort comprised 39 patients (stool samples) with UC stratified by 5-ASA use: current (n = 20), prior (n = 15), and non-users (n = 4) (Fig. 3b)^36^. Among 20 subjects with UC reporting current 5-ASA use, 5-ASA was detectable in 18 individuals. Cholyl–5-ASA was detected in all of them (100%). Of the 15 individuals reporting prior use of 5-ASA, 6 had detectable levels of 5-ASA. Cholyl–5-ASA was detected in 3 of these 6 individuals (50%), consistent with reduced detection after discontinuation, and 2 of 4 reported as non-users. The odds ratio comparing current users to prior and non-users was 22.5 (95% CI: 3.5–268, Fisher’s exact p = 6.86×10^-5^). Notably, two individuals labeled as non-user in their metadata had detectable cholyl–5-ASA, and in one of them, we detected 5-ASA itself as well (Supplementary Fig. S1b). Correcting based on detection of the drug and metabolites with respect to these two cases increased the odds ratio to 29 (95% CI: 4.36–361, Fisher’s exact p = 0.000018).

In a third, paediatric cohort of 54 children (28 healthy controls, 26 diagnosed with very-early onset IBD, age 0.5–5), 10 individuals were receiving 5-ASA or 5-ASA prodrugs: two as balsalazide, five as sulfasalazine, and three as mesalazine (5-ASA)^39^. Among these 10 individuals, 5-ASA was detectable in 8 individuals (Supplementary Fig. S1c), and cholyl-5-ASA was detected in 9 individuals (Fig. 3c).

The fourth data set available was a surgery IBD intervention cohort that had samples from 34 UC and 24 CD volunteers, often multiple samples per individual. Among the 30 subjects with UC or CD reporting 5-ASA use, 5-ASA was detectable in 25 individuals (21 UC and 4 CD).

Cholyl–5ASA was detected in 10 of these 25 individuals with detectable 5-ASA (40%), including 9 with UC and 1 with CD. It showed that cholyl–5-ASA is primarily, but not exclusively, observed in UC when the use of 5-ASA medications is documented (Fig. 3d)^40^. As the goal of the study was to monitor outcome of the surgery, the use of 5-ASA was not always specifically documented.

To further validate the presence of this drug conjugate in subjects treated with 5-ASA or associated prodrugs in people outside the US, we examined one additional CD cohorts from the United Kingdom: the iPENS cohort (individuals [n = 48]; stool samples [n = 214]).

We observed cholyl-5-ASA present in stool samples collected from patients with CD in the first IBD cohort mentioned above, it should be noted that the use of 5-ASA medications in CD is common but also controversial^41^. However, these observations prompted us to question if bile acid–5-ASA conjugates are commonly detected in CD data as well and explored the iPENS CD study that included 14 participants reporting the use of 5-ASA medications^41,42^. In the iPENS data, 5-ASA was detected in 9/14 individuals, in which all 9 reported current 5-ASA use (100%). It was not detected from 12 participants (39 longitudinal samples) that did not self-report 5-ASA treatment for which we had complete metadata for prescribed medications. Cholyl–5-ASA was detected in the same 9 individuals with CD (100%) (Fig. 3e). These 9 patients provided a total of 39 longitudinal stool samples where cholyl–5-ASA was detected.

We also set out to assess whether the relative 5-ASA levels dictate the amount of bile acid–5-ASA formed. If conjugation were driven primarily by substrate availability, a monotonic relationship between parent drug abundance and conjugate levels would be expected. We therefore examined the relationship between 5-ASA and distinct classes of bile acid–5-ASA conjugates and observed positive correlations across adult IBD cohorts (Spearman’s ⍴ ranging from 0.53-0.89), whereas correlation was weaker and not statistically significant in the paediatric cohort (Spearman’s ⍴ ranging from 0.21-0.55; Supplementary Figure S3). This suggests that while 5-ASA levels improve the likelihood the bile acid conjugates are observed but are not the sole driver and that there are other factors such as the availability of an “activated” cholic acid and the enzymes, possibly microbial derived, able to process this transformation will likely also be critical determinants.

To quantify cholyl–5-ASA levels, we prospectively enrolled a sixth IBD cohort consisting of 10 age-matched individuals without an active 5-ASA prescription and 10 individuals with an active 5-ASA prescription. For the patients prescribed 5-ASA, 2 had UC and 8 had CD. In the comparison (non-5-ASA cohort), two had UC and eight had CD. Of the 10 individuals with an active 5-ASA prescription, the drug itself was not detected in three participants (Supplementary Fig. S1g). Among the seven individuals in whom 5-ASA was detected, concentrations ranged from 1 µM to 9 mM. Targeted LC-MS quantification detected the cholyl–5-ASA conjugate in the same seven individuals (100%), with concentrations ranging from 0.02 to 48 µM (Fig. 3g).

While all the results from the reanalysis of the five public datasets and the quantification data are correlative, they do not establish its effect on activity. Instead, they suggest an intriguing hypothesis: 5-ASA conjugation to bile acids - potentially mediated by microbiota - may influence the drug’s therapeutic efficacy. If these bile acid–5-ASA conjugates alter 5-ASA’s anti-inflammatory properties then differences in microbial composition or lower microbial biomass^43^ could help explain differences in treatment response seen with 5-ASA medications.

### Human gut bacteria can conjugate 5-ASA to bile acids

As the microbiome is intimately linked to IBD^43–47^, as well as 5-ASA activity^48–50^ and we discovered that microbes can carry out the conjugation of bile acid reactions^11^, there was a reasonable chance that microbiota could catalyze this transformation, however the specific bacteria responsible for producing bile acid–5-ASA conjugates is currently unknown. To gain insight into which taxa may be capable of catalyzing bile-acid–5-ASA conjugation, we reprocessed a dataset, for which both metagenomics and untargeted metabolomics data were acquired^36^, implementing the latest host read removals and genome coverage filter^51,52^. After subsetting this dataset for subjects with detectable levels of fecal 5-ASA (n = 53) to focus on individuals with substrate availability, we integrated metagenomics and metabolomics data using joint-RPCA^53^. This process identified many bacterial species as positively associated with cholyl–5-ASA, belonging to the *Bifidobacterium*, *Lactobacillus*, *Collinsella*, *Actinomyces*, *Streptococcus*, *Klebsiella*, *Clostridium*, and *Enterococcus* genera (Fig 4a). This observation motivated us to test an array of bacteria to assess if they could generate cholyl–5-ASA and other drug–bile acid conjugates in cultures.

**Figure 4.**
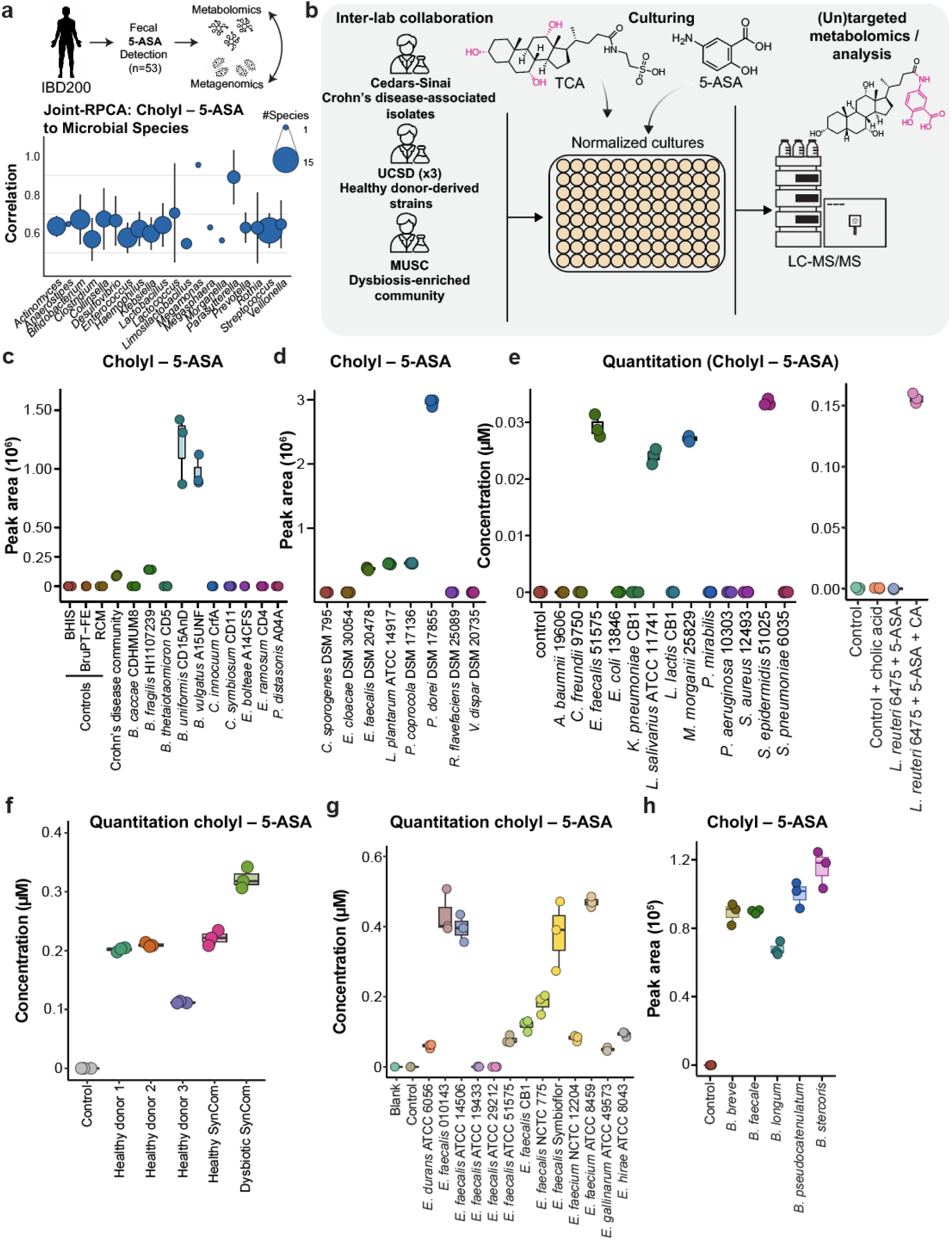
Human gut bacteria can conjugate 5-ASA to cholic acid. a, Joint-RPCA analysis of metagenomics and metabolomics data acquired in the IBD200 cohort. Overview at Genus level of selected OGUs positively correlating with cholyl–5-ASA. Balloon plot display mean correlation values + standard deviation of selected OGUs of interest collapsed at Genus level. Only positive correlation values > 0.5 were taken into consideration (higher values indicate stronger correlation). Sizes of the dots reflect the number of collapsed significant species per genus. **b,** Overview of experimental workflow: three independent laboratories tested the ability of gut bacteria to produce bile acid–5-ASA conjugates by adding bile acids and 5-ASA to growth media. Absolute quantification was performed in one lab, while relative quantifications were obtained in the others. **c,** Cholyl–5-ASA production in 11 single microbial cultures derived from mucosal isolates of individuals with IBD after 72 h. No production is observed at baseline (0 h). **d,** Production of cholyl–5-ASA by eight tested strains in a second laboratory after 72 h. No production is observed at baseline. **e,** Production of cholyl–5-ASA in 15 strains after 72 h. Here the controls are no culture added. These were done as two separate experiments hence they are shown as two separate panels. (b) and (c) represent relative while (d) represent absolute quantification from two different experiments in the same lab. Each experiment in all labs was carried out as three biological replicates. **f,** Quantitation (µM) of cholyl–5-ASA of cultures of microbial communities obtained from three human healthy donors as well as two microbial consortia. **g,** Quantitation (µM) of cholyl–5-ASA across a range of *Enterococcus* species. **h,** Production of cholyl–5-ASA in five *Bifidobacterium* species isolated from humans after 72 h. No production is observed in the medium supplemented with the substrates.

To this end, we collaborated with five different laboratories that have ongoing culturing experiments of bacterial species from human microbiota. This unbiased approach allowed us to assess a broader ability of gut bacteria to produce bile acid–5-ASA conjugates when bile acids and 5-ASA are separately added to the growth medium by experts familiar with growing these microbes (Fig. 4b). We are reporting on cholyl–5-ASA for all the studies as all studies had the analysis of cholyl–5-ASA in common. One group performed absolute quantitation, while the others provided relative quantifications. In total, seven different culturing experiments were performed. Collectively, all experiments demonstrated that several human gut bacteria harbor the genomic machinery to produce bile acid–5-ASA conjugates.

In the first lab’s initial experiment, we evaluated mucosal bacterial isolates obtained from individuals with CD. Three of the 10 single-strain cultures produced cholyl–5-ASA, and all belonged to the genus *Bacteroides*, with *B. vulgatus* and *B. uniformis* showing the highest levels (Fig. 4c). In the second lab, eight additional strains were tested, and four produced cholyl–5-ASA; the most prolific producer was *Phocaeicola dorei* (formerly *Bacteroides dorei*; Fig. 4d). The third and fourth experiments quantified production across 11 strains, which generated cholyl–5-ASA at concentrations of 0.03 µM and 0.15 µM, respectively (Fig. 4e). Across all four studies, *E. faecalis* was consistently identified as a producer, although the production appeared to be strain-specific.

We next asked whether human microbial synthetic communities, rather than individual isolates, could generate cholyl–5-ASA. All three undefined communities and both defined consortia produced the compound, though at varying levels (Fig. 4f). Because *E. faecalis* appeared as a producer in every preceding experiment, we investigated strain-level variation within this species. Not all *Enterococcus* species - or even all *E. faecalis* strains - encode bile salt hydrolase genes, which are required for bile-acid amino-acid amidation as previously shown^54,55^. We therefore tested ten *E. faecalis* strains and three phylogenetically related strains (Fig. 4g). We found that only the *E. faecalis* strains with a bile salt hydrolase could make the cholyl–5-ASA. Finally, we tested five *Bifidobacterium* species isolated from human volunteers for their ability to conjugate 5-ASA to bile acids. All five species produced cholyl–5-ASA, showing conserved conjugation capacity within this genus (Fig. 4h).

To summarize these seven culture-based experiments, many gut bacterial taxa appear to be able to produce cholyl–5-ASA. Within the phylum *Bacteroidota*, *Bacteroides* species (e.g., *B. fragilis, B. uniformis, B. vulgatus*) appeared to be key producers. Among *Bacillota*, species of *Lactobacillus* and *Enterococcus* also exhibit activity. Other phyla, including *Pseudomonadota* and *Actinomycetota* (e.g., *Bifidobacterium* species), contribute minimally or not at all, indicating that cholyl–5-ASA synthesis is primarily carried out by specific members of *Actinomycetota*, *Bacillota*, and *Bacteroidota*. This taxonomic distribution is also largely consistent with the availability of the bile salt hydrolase gene in the genome by producers that we, and others, recently found to have bile acid transamination activity^54,55^.

### Bile salt hydrolases conjugate 5-ASA to bile acids

Most of the organisms that we show to be able convert taurocholic acid to cholyl**–**5-ASA have one bile salt hydrolase gene (*bsh*) in their genome, and some have multiple. We and others have recently demonstrated that *bsh* genes encode for bifunctional enzymes^54–56^. These enzymes hydrolyze taurine-and glycine-conjugated bile acids, as was established in the 1960s, but also catalyze the transamination reaction of taurine conjugated bile acids where the taurine is replaced with amine containing substrates, including 5-ASA. In addition, there are organisms that can carry out transaminase activity that have unrecognizable bile salt hydrolases. BSH from different bacteria have different substrate specificities^57^. They have different transamination specificities for both amines and taurine activated bile acids. To test whether BSH activity is required for the formation of bile acid**–**5-ASA conjugates, and if any of them can catalyze the transamination with 5-ASA, we tested nine engineered *E.coli* constructs in which different *bsh* genes were integrated in the genome^17,58^. The introduced *bsh* genes originated from *B. vulgatus, D. newyorkensis, L. gasseri, B. fragilis, B. uniformis, E. plexicaudatum, Lachnospiraceae CAG-95.* As controls, *E.coli* expressing the green fluorescent protein (GFP) instead of a *bsh* gene was included as well as a control. Engineered *E. coli* strains were incubated with taurocholic acid, taurodeoxycholic acid, and taurolithocholic acid with 5-ASA and sampled at 0 h and 48 h (Fig. 5). Five of the nine engineered strains are active as evidenced by the depletion of the taurine bile acid conjugates (Fig. 5a). For the remaining engineered strains, absence of detectable activity cannot be unambiguously attributed to lack of expression, protein misfolding, or divergent substrate specificity based on metabolite measurements alone. The different *bsh* engineered *E.coli*’s that we found to be able to delete taurine conjugates all displayed different enzyme-specific substrate preferences for production of 5-ASA conjugates but also of amino acids and polyamines (Fig. 5b). Consistent with observations in bacterial cultures, expression of BSH1 from *B. vulgatus* in *E. coli* was the only construct that produced all three bile acid amidates, including cholyl–5-ASA, whereas the BSH2 from *B. vulgatus* did not produce cholyl–5ASA. Instead, *B. vulgatus* BSH2 preferentially produced deoxycholyl**–**5-ASA and lithocholyl**–**5-ASA, showcasing complementary substrate specificity between BSH homologs. Lithocholyl**–**5-ASA was also detected across multiple BSHs such as *D. newyorkensis* BSH1, *L. gasseri*, and *Lachnospiraceae* CAG-95, providing direct evidence that microbiome-derived BSHs can indeed produce the bile acid–5-ASA conjugates. But that multiple BSH genes, originating from different organisms, reinforces that many, but not all, organisms that carry a *bsh* gene are able to form 5-ASA conjugates with bile acids. Prior studies have shown that 5-ASA fails to suppress inflammation in the absence of a microbiome suggesting that microbial metabolism may be required for activity. Together with our observation that 5-ASA is conjugated by microbiota, more specifically the BSH enzyme, these findings raise the possibility that bile acid**–**5-ASA conjugation alters the biological activity of 5-ASA.

**Figure 5.**
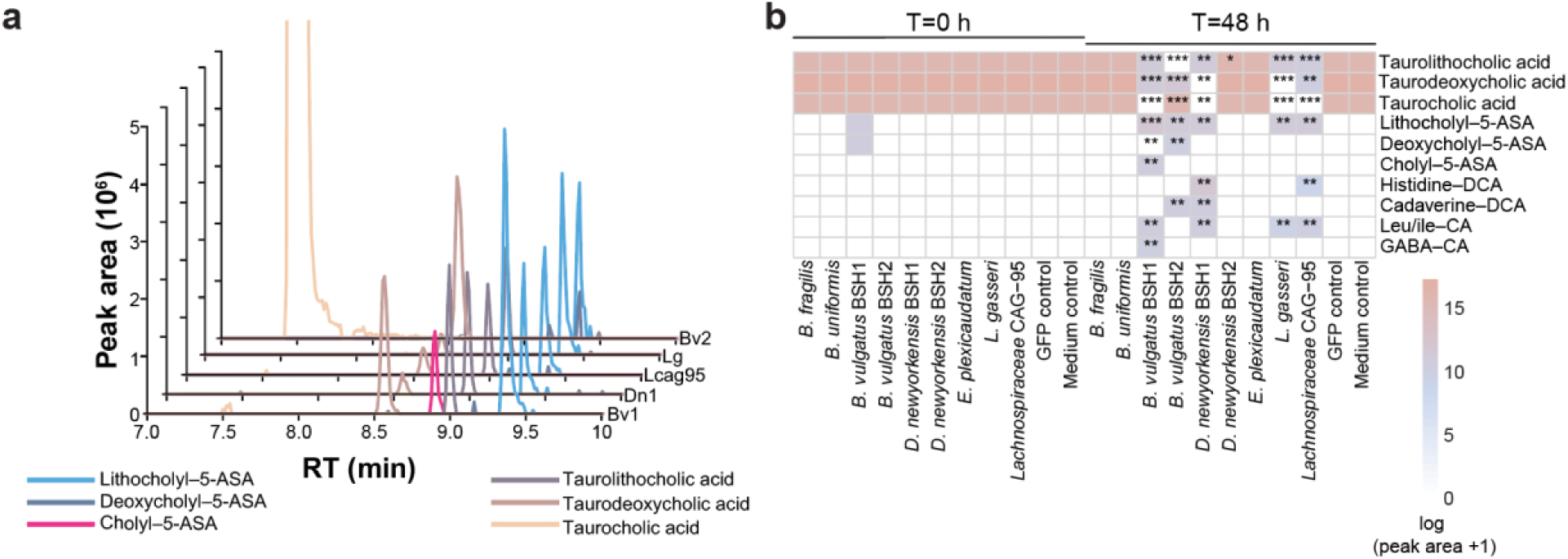
Distinct bile salt hydrolases selectively catalyze bile acid–5-ASA conjugations from taurine conjugates of lithocholic acid, deoxycholic acid, and cholic acid. a, Representative extracted ion chromatograms of lithocholyl–5-ASA (*m/z* 512.3373; light blue), deoxycholyl–5-ASA (*m/z* 528.3342; dark blue), cholyl–5-ASA (*m/z* 544.3269; pink), taurolithocholic acid (*m/z* 484.3091; purple), taurodeoxycholic acid (*m/z* 500.3040; brown), and taurocholic acid (*m/z* 516.2990; orange) at 48 h after culturing for the five engineered constructs that showed activity. **b,** Heatmap showing the median-log transformed peak area of bile acid conjugates detected in *E. coli* expressing *bsh* genes of gut microbes as well as a GFP control. Color intensity represents the median of three biological replicates. Statistical analysis was performed using a two-sided unpaired t-test. Significance levels are indicated on the heatmap as follows: *p* < 0.05 (*), *p* < 0.01 (**), *p* < 0.001 (***), and *p* < 0.0001 (****). The *B. vulgatus* BSH1 is very efficient and by the time the samples were frozen ∼1-10 min after addition of the microbial culture to stop the reaction partial (∼1-2%) conversion was already observed. Statistical analysis was performed using t-test. Bv1, *B. vulgatus* BSH1; Dn1, *D. newyorkensis* BSH1; Lcag95, *Lachnospiraceae* CAG-95; Lg, *L. gasseri*; Bv2, *B. vulgatus* BSH2; 5-ASA, 5-aminosalicylic acid; DCA, deoxycholic acid; CA, cholic acid.

### Unexpected enhancement of activity of 5-ASA by bile acid conjugation

We wanted to understand if bile acid–5-ASA conjugation affected 5-ASA activity. *In vivo*, 5-ASA is known to affect PPAR-γ^59,60^ and to influence multiple CD4^+^ T cell subsets, including Foxp3^+^ regulatory T cells^61^, and IFN-γ producing Th1 cells^62^ reducing the severity of DSS induction of colitis in animal models^59,60^. As we do not have such assays in our lab, we collaborated with three different labs that routinely carry out relevant assays to assess the effect of bile acid conjugation to 5-ASA (Fig. 6a). The effect of 5-ASA compared to cholyl–5-ASA on PPAR-γ and CD4^+^ T cells responses were measured in cell-based assays. In parallel, the capacity of each compound to suppress DSS-induced colitis was evaluated *in vivo*. Chemical modification of small molecule drugs often leads to partial or complete loss of activity; however, contrary to this expectation, cholyl–5-ASA consistently exhibited greater activity than unconjugated 5-ASA across all three independent assay systems that were evaluated (Fig. 6).

**Figure 6.**
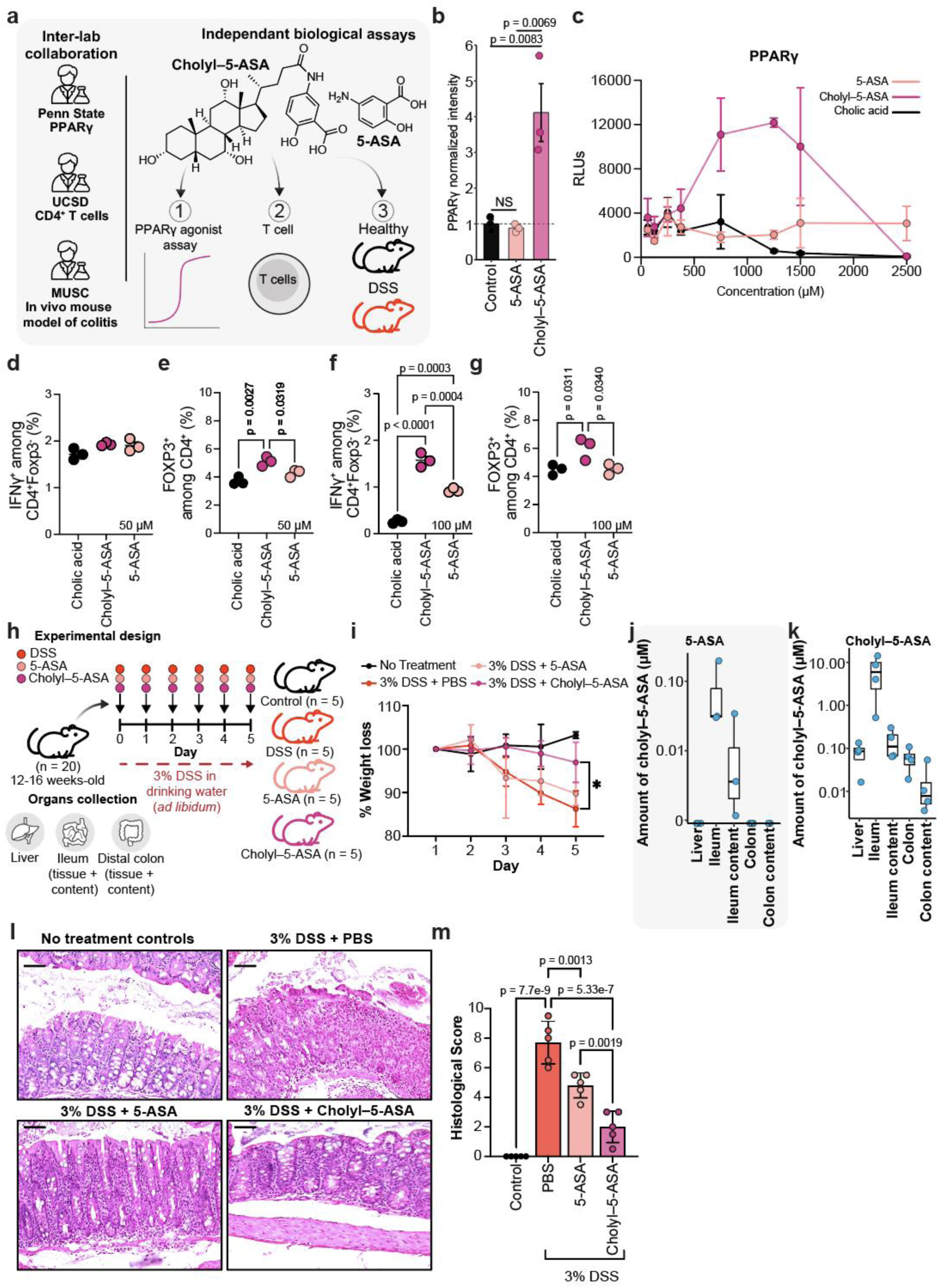
Bile acid conjugation enhances 5-ASA activity. a, Overview of experimental workflow to get biological activity insights: three independent laboratories (Penn State, UCSD, and MUSC) tested two cell-based assays while the third one tested the ability of cholyl–5-ASA compared to 5-ASA in a DSS-colitis mouse model. **b,** PPAR-γ reporter activity in human cells treated with 5-ASA and cholyl–5-ASA at 1 mM, 10X concentration of cholyl-5-ASA where saturation of activity is observed. Values are normalized to control (value = 1). Statistical analysis was performed using one-way ANOVA with Tukey’s post-hoc test after confirming normality (Shapiro-Wilk) and homogeneity of variance (Levene’s test). **c,** PPAR-γ transcriptional activity was measured using a reporter and is shown as relative luminescence units (RLUs) across increasing concentrations of 5-ASA, cholyl–5-ASA, and cholic acid. Data points represent the mean values with error bars indicating the standard deviation. **d, f,** Frequency of IFN-γ^+^ among CD4^+^Foxp3^-^ T cells and **e, g,** Frequency of Foxp3^+^ cells among CD4^+^ T cells isolated from C57BL/6J WT following treatment with cholic acid, cholyl–5-ASA, and 5-ASA at 50 µM (d, e) and 100 µM (f, g). Statistical significance was assessed using one-way ANOVA with Tukey’s post-hoc test. **h,** Experimental design of the DSS-colitis mouse model. **i,** Effect on mouse weight in 3% DSS treated mice against 5-ASA and choly–5-ASA treatment. Two-way ANOVA was used to evaluate significance. **j,** Quantitation of cholyl-5-ASA across organs and contents in mice gavage with 5-ASA and **k,** cholyl–5-ASA. **l,** Representative images of hematoxylin and eosin (H&E) stained of formalin-fixed paraffin-embedded colon sections and **m,** their histological score. Statistical analyses were performed using one-way ANOVA with Tukey’s post-hoc test. Exact *P* values are indicated where applicable. GraphPad Prism (version 10.6.1) was used for selected data visualization and statistical analyses.

The first assay evaluated was a cell-based PPAR-γ assay. Although prior studies have shown that 5-ASA can activate PPAR-γ in IBD models *in vivo*^59,60^, they did not determine whether this activation is mediated by 5-ASA itself or bile acid–5-ASA as they were not aware of the existence of bile acid–5-ASA conjugates produced by gut microbiota. Remarkably, cholyl–5-ASA is much more effective and potent than 5-ASA in inducing PPAR-γ activation compared to background control although at higher concentration, and well above the observed concentrations found in humans, it does appear to become toxic to cells (Fig. 6b,c).

The second assay evaluated was a cell-based T cells differentiation assay. To test whether cholyl–5-ASA modulates T cells function, we incubated T cells with cholyl–5-ASA, 5-ASA, and cholic acid. We also found that cholyl–5-ASA elicited a stronger response in CD4⁺ T cells than 5-ASA, characterized by increased Foxp3 expression, as well as IFN-γ induction (Fig. 6d-g). These findings support a model in which microbial conversion of 5-ASA to cholyl–5-ASA modulates CD4⁺ T cell responses.

In the final assay, we monitored the effects of 5-ASA and cholyl–5-ASA in a 3% DSS-induced colitis mouse model in specific pathogen-free mice (Fig. 6h). This experiment mimics maintenance of remission in people rather than reversal of inflammation after onset, as DSS was introduced together with 5-ASA or cholyl–5-ASA. Assessment of weight loss has been shown to reflect the degree of outcome of interventions in a DSS model and is one of the most reliable and cost effective measures in this model^63^. We therefore measured the weight of the animals in each of the interventions. We found that mice treated with DSS or DSS + 5-ASA had lost ∼10% of their weight by day 5. In contrast, mice treated with DSS + cholyl–5-ASA exhibited little weight loss, more comparable to untreated non-DSS controls (Fig. 6i). Quantitation of cholyl–5-ASA in the ileum and colon contents, the ileal and colonic tissue, and liver, from the DSS treated mice revealed that cholyl–5-ASA is formed in the 5-ASA treated mice at ∼100 fold lower amounts compared to direct administration of cholyl–5-ASA (Fig. 6j,k). Specifically, cholyl–5-ASA was detected at 10 µM in the ileal content when it was directly administered, and at 0.1 µM when 5-ASA was provided instead. To assess intestinal inflammation, a histological colitis score was generated as previously described^64,65^ (Fig. 6m). As expected, DSS-treated mice exhibited high levels of colonic inflammation, characterized by immune cell infiltration, marked reduction of mucus-producing goblet cells (Fig. 6l). Administration of 5-ASA significantly reduced colonic inflammation by approximately 50% (mean score: PBS, 7.7). Cholyl–5-ASA induced a much stronger effect than 5-ASA (mean score: 5-ASA, 4.8; cholyl–5-ASA, 2.0; *p*-value = 0.0019), reducing inflammation by about 75% compared to DDS-treated mice, restoring goblet cells (Fig. 6l,m).

## Discussion

Pan-repository scale of untargeted metabolomics analysis has only recently become feasible^26^. Using this capability, we identified six unique MS/MS that corresponded to specific bile acids found exclusively in RA and IBD patients. Here, we report the unexpected but data science driven discovery of microbial bile acid–drug conjugates extending our understanding of bile acid metabolism together with a drug beyond conventional pathways.

This is the first report of bile acid–5-ASA detection in humans and animals after 5-ASA administration, a very old group of medications. Since its approval for human use in 1950, recognition that 5-ASA in 1977 was the active form of sulfasalazine^33^ for the treatment of ulcerative colitis, and subsequent introduction of its intestinal deliverable version and prodrug versions in the 1980s and 1990s, 5-ASA has, even with the introduction of biologics, due to low cost and fail first insurance practices^66^, has remained a cornerstone treatment as maintenance therapy. Despite its introduction over 75 years ago, its mechanism of action is, surprisingly, still rather ill-defined^67^. A systematic review of 39 different studies reported that there is no explanation as to why levels of 5-ASA are not linked to therapeutic efficacy^68^. Although 5-ASA is proposed to act through multiple linked mechanisms, including regulation of cyclooxygenase expression and lipoxygenase pathways, scavenging reactive oxygen species, suppression of NF-κB signaling, activation of PPAR-γ, and modulation of immune cell infiltration^69,70^, none of these mechanisms explained why a high amount of 5-ASA - often reaching well over tens of mM does not result in a therapeutic response in all individuals.

Analysis of *in vitro* microbial cultures demonstrated that specific gut bacteria can conjugate 5-ASA to bile acids, producing cholyl–5-ASA and related derivatives. Direct administration of cholyl–5-ASA-made by microbes-in a DSS colitis mouse model conferred stronger suppression of DSS induced colitis than 5-ASA alone. It is even plausible that cholyl–5-ASA is the active form of the drug that drives decreases in inflammation, and not 5-ASA, as previous literature found that the microbiome is essential for the activity of 5-ASA in a DSS colitis model^71,72^. That microbiota affects 5-ASA therapy is also observed in fecal microbiota transplantation (FMT) and probiotic studies in humans and animals. Co-administration of probiotics *Bifidobacterium* and mixed *Lactobacillus/Bifidobacterium* with 5-ASA in 13 studies increased the efficacy of 5-ASA therapy^73^. Notably, the probiotic formulations tested in those 13 studies are primarily *Bifidobacterium* and mixed *Lactobacillus/Bifidobacterium* preparations. These two genera commonly encode BSH, and the particular species used in the preparations are ones that we show here can produce bile acid-5-ASA conjugates. Similarly, add-on therapy of a fecal transplant in a randomized human clinical trial resulted in a statistically significant added benefit to 5-ASA treatment and, in a separate study, also trended towards improved outcomes of 5-ASA treatment, but was underpowered^74,75^. Animal studies also report the combined effect of 5-ASA and microbes. For instance, co-administration of *Lactobacillus casei* - a bacterium carrying *bsh* genes - with 5-ASA (mesalazine) has been shown to increase the drug’s effectiveness in an animal model of ulcerative colitis^74^. Together, these observations are consistent with a microbially derived 5-ASA metabolite being the active form of the drug that drives the reduction of inflammation. Perhaps the lack of medication response in many patients is due to a combination of both microbial composition, which is variable in IBD^76,77^, and microbial loads^43^.

Once we found the bile acid–5-ASA conjugates to be more active in colitis models, we explored the literature about what is known about bile acids and 5-ASA. These previous studies include studying co-administration of bile acids with 5-ASA^75,78^, as well as chemically linking bile acids to 5-ASA to improve therapeutic delivery. When bile acids were co-administered, the activity of 5-ASA improved. Specifically, ursodeoxycholic acid (UDCA), another clinically approved drug, together with 5-ASA was shown to have enhanced anti-inflammatory effects^78^. Other studies focused on synthesizing UDCA–5-ASA and chenodeoxycholyl–5-ASA (CDCA–5-ASA) conjugates as colon-targeted prodrugs^76,77,79^. In guinea pig colitis models, ursodeoxycholyl–5-ASA and chenodeoxycholyl–5-ASA conjugates produced strong reductions in ulceration and bleeding^77^, and improvement in body weight, similar to the effects we observe in preventing colitis mice treated with cholyl–5-ASA in this study. These conjugates were designed in those studies to pass through the upper digestive tract and release free 5-ASA once in the colon. The Guinea-Pig study and corresponding patent never considered if the bile acid–5-ASA conjugates could be active themselves as they regarded these conjugates primarily as delivery vehicles. This prior literature did not know nor consider that similar bile acid conjugates are actively produced in humans that are taking 5-ASA and its prodrugs - which we show here to be the case.

Our observations, alongside prior research, suggest that 5-ASA is largely inactive in its parent form and must be converted by the microbiota into active metabolites to exert its effects (Fig. 7). Mechanistically, cholyl–5-ASA exhibits enhanced PPAR-γ activation and modulates T cell expression of IFN-γ among CD4^+^Foxp3^-^ cells and induction of Foxp3^+^ T cells, providing a plausible explanation for its superior anti-inflammatory profile compared to 5-ASA alone. These results resolve a paradox where 5-ASA fails to suppress colitis in the absence of a microbiome: without microbial metabolism, its most potent forms are never generated. Together, these findings reframe 5-ASA pharmacology as a microbial–host co-metabolic process rather than the isolated action of the parent drug. Given that 5-ASA concentrations can reach levels exceeding 10 mM, a conversion rate of only 0.01% would be sufficient to achieve the 10 µM of cholyl–5-ASA observed to suppress injury in our *in vivo* DSS study (Fig. 6). Quantitative human data show that cholyl–5-ASA levels can reach up to 9.7% of measured 5-ASA levels. While this may be influenced by differential pharmacokinetics, it suggests that conversion rates significantly higher than 0.01% are likely (Figure 3d, Supplementary Table S1). Consequently, bile acid–5-ASA conjugates are formed in sufficient quantities to account for the drug’s primary immunomodulatory activity.

**Figure 7.**
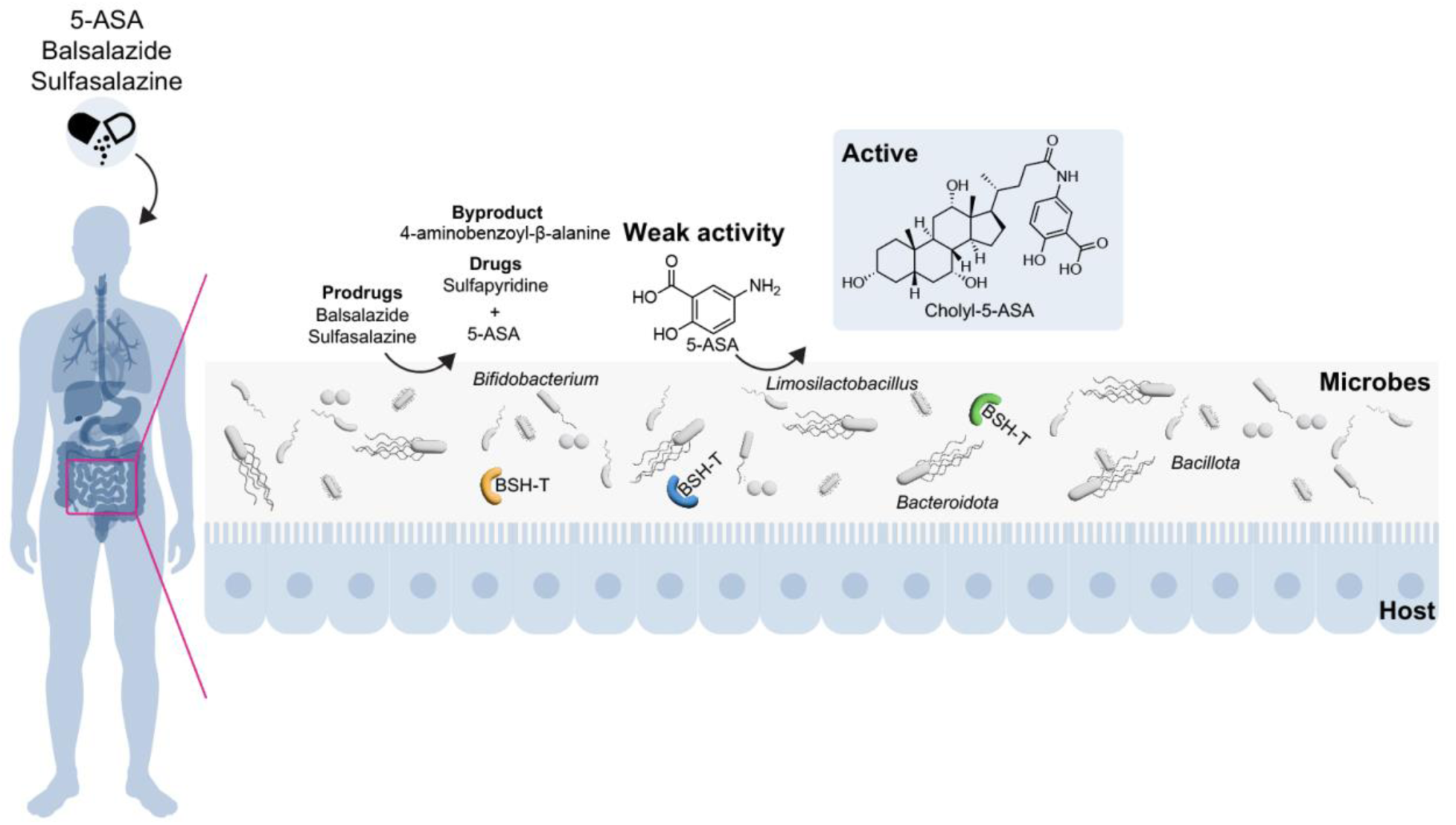
Overview of 5-ASA metabolism and its relationship to bile acid conjugation and activity. Current understanding of metabolism and how it affects activity of 5-ASA medications for maintenance of remission. Following oral or rectal administration, 5-ASA and its prodrugs (e.g., sulfasalazine, balsalazide) enter the gut lumen, where they undergo conversion to bile acid conjugates. BSH-T, bile salt hydrolase - transaminase.

Beyond this specific drug, our study establishes a unique functional role for microbial bile acids in drug metabolism and illustrates how pan-repository metabolomics can reveal previously unrecognized microbiome-dependent biochemical transformations with direct therapeutic relevance. In this context, cholyl–5-ASA serves both as a mechanistic biomarker of 5-ASA activity and as a proof-of-concept for bile acid mediated therapeutic optimization.

More broadly, they support an emerging yin and yang framework of microbiome-mediated drug metabolism, in which microbial transformations are essential for drug efficacy yet can either potentiate or inactivate therapeutics. Within this framework, microbial bile acid conjugation emerges as a previously unrecognized biochemical function that directly shapes therapeutic activity. Consistent with this model, direct administration of bile acid–5-ASA conjugates enhanced efficacy in animal experiments, suggesting a potential strategy to bypass microbiome-dependent non-responsiveness.

### Limitations, future research and potential translational implications

Despite these insights our study provides into bile acid-mediated drug activation by microbes, several limitations remain. While our data support bile acid conjugation as an activating step for 5-ASA in rodents, evidence in humans is currently correlative. Establishing clinical relevance, particularly in maintenance therapy rather than acute inflammation, will require controlled prospective studies. Formal evaluation of the safety, dosing, and pharmacokinetics of bile acid–5-ASA conjugates will necessitate dedicated preclinical and early-phase clinical trials beyond the scope of this work.

Interindividual variation in microbiota composition is likely to influence bile acid–5-ASA formation, but the contributions of diet, medications, host genetics, bile acid pool composition, and disease localization remain unresolved. Although cholyl–5-ASA is detectable systemically, it is unclear whether bile acid conjugation primarily enhances systemic exposure, prolongs local intestinal activity, or both. Addressing these questions will require comprehensive pharmacokinetic studies across diverse microbial states, including germ-free and antibiotic-treated models.

An important open question is whether bile acid–5-ASA conjugates constitute major mediators of therapeutic efficacy in humans. This hypothesis is supported by the microbiome dependence of 5-ASA activity in animal models and by reduced efficacy in severe IBD, where key microbiota are depleted. Large-scale clinical studies correlating bile acid–5-ASA abundance with treatment response, including under antibiotic exposure, could establish these conjugates as predictive biomarkers.

If validated, this mechanism suggests multiple translational opportunities, including personalized diagnostics, microbiome-or enzyme-based adjunct therapies, and the development of bile acid-inspired prodrugs.

## Methods

### Biospecimen sample processing

A prospectively enrolled IBD cohort consisting of 20 patients (10 patients under 5-ASA / 10 aged-matched IBD controls without 5-ASA prescription) was used for quantification (IRB#202017; Vanderbilt University Medical Center). Cholyl–5ASA was also quantified from bacterial extracts. Human and murine feces were weighed on an electrical scale (Ohaus Corporation, Parsippany, NJ) and approximately ∼150 mg of the feces were added to 2 mL fast prep-tubes harboring 0.1 g of Lysing Matrix D 1.4 mm ceramic beads. To each tube, ice-cold ethanol was added to generate a fecal dispersion at a density of 250 mg/mL^80^. Murine liver was also weighed and approximately∼150 mg of the liver was added to 2 mL fast prep-tubes that each contained 0.1 g of 1.4 mm ceramic beads. All tubes containing either fecal or liver tissues were homogenized on a Benchmark “Beadbug” 6 Microtube Homogenizer (Catalog #BS-BEBU-6; Stellar Scientific, Baltimore, MD, USA) at 4.0 m/s for 20 s for two cycles. Tubes were centrifuged at 12,000 x *g* for 5 min to pellet any solids and supernatant was transferred to new tubes for storage at-80 °C until bioanalysis could be performed. Prior to 5-ASA bioanalysis, each specimen was diluted 20-fold in a solution consisting of acetonitrile: water (9:1, *v:v*) that contained 0.1% formic acid. For cholyl–5-ASA bioanalysis, each specimen was diluted 20-fold in a solution consisting of methanol: water (1:1, *v:v*). In each instance, a 10 µL volume of sample was injected onto the liquid-chromatography high-resolution mass spectrometer system.

### Reanalysis of public dataset from GNPS/MassIVE

To evaluate the prevalence of the drug-bile acid conjugate in other datasets, we used FASST to find datasets available in the public domain in which the cholyl–5-ASA conjugate was found. The search resulted in finding several datasets in which cholyl–5-ASA was detected in GNPS/MassIVE. In addition to the RA dataset, four other datasets were selected for reanalysis:

(1) a study on IBD patients (MSV000084775; fecal samples), (2) another cohort of IBD patient (MSV000082094; fecal samples), (3) a paediatric IBD cohort (MSV000097610; fecal samples) and (4) a longitudinal surgery IBD cohort (MSV000082221; fecal samples). For each dataset, mzML files were downloaded from GNPS/MassIVE and processed using MZmine (parameters and versions used are detailed in Supplementary Table S2). The output from MZmine, the csv file containing the peak areas and the mgf file with the MS/MS spectral information for each feature were imported into GNPS2 to perform the feature-based molecular networking (FBMN) workflow. The parameters used in FBMN were the same for all datasets in which the precursor and fragment ion tolerance were set to 0.02, no filtering steps, and for the networking and the library search, the minimum cosine was set to 0.7 with a number of matched peaks of 6. The FBMN job links for each dataset can be accessed here:

Rheumatoid arthritis cohort^29^ (MSV000084556): https://gnps2.org/status?task=dfdf2c8d87d544dfa7eb5f49d6711112

Inflammatory bowel disease cohort 1^36^ (MSV000084775): https://gnps2.org/status?task=53b4e6f2a0234e18a6e036b7a5116582

Inflammatory bowel disease cohort 2^36^ (MSV000082094): https://gnps2.org/status?task=1999554e1d514150b0ac9fb6882937f7

Paediatric IBD cohort 3^39^ (MSV000097610): https://gnps2.org/status?task=4559be69e9984f60a36d47f1025f145f

Inflammatory bowel disease cohort 4^40^ (MSV000082221): https://gnps2.org/status?task=afe5f5aa8b2945f0a3c4a53c9e5e61f4

### Repository-scale analysis using FASST

FASST, a fast version of MASST that searches any MS/MS spectra against the four main metabolomics repositories GNPS/MassIVE, Metabolights, Metabolomics Workbench, and NORMAN, was used through the GNPS2 FASST batch workflow available here https://gnps2.org/homepage. A mgf file containing a list of MS/MS spectra of all the candidate bile acids (accessible here https://external.gnps2.org/gnpslibrary) was used as input in the FASST search. The parameters were the following: minimum cosine of 0.7, precursor and fragment ion tolerance of 0.02. The Pan-ReDU metadata (https://redu.gnps2.org/selection/) (August, 2025) was downloaded and imported into RStudio (version 4.5.2) as well as the output from the FASST search for downstream analysis. The reverse metabolomics workflow was used to associate each candidate bile acid retrieved from the FASST search with human disease phenotype (using the DOIDCommonName column from the Pan-ReDU metadata). Only human-derived entries were retained for the analysis. All matches were filtered for a cosine similarity score of at least 0.7, with at least 4 matching peaks (typically resulting in a false discovery rate of < 1%)^81^. To reduce spectral redundancy and downstream analysis, neutral water losses were excluded and only bile acid candidates observed in at least two different health phenotypes were retained. The final matrix, based on presence and absence of the candidate bile acids, was summarized and visualized using the ComplexUpset package (version 1.3.3)^82^. The chord diagram was generated using the package circlize (version 0.4.16) available on CRAN^83^.

### Bacterial strains, culture conditions, and extraction

Bacterial strains used in this study are listed in Supplementary Table S3. Bacteria were started from glycerol stock and incubated in filtered brain-heart infusion (BHI) in an anaerobic chamber (20% CO_2_, 5% H_2_, and 75% N_2_). Details on the medium composition can be found in Supplementary Table S4. Bile acids (taurolithocholic acid; TLCA, taurocholic acid; TCA, taurodeoxycholic acid; TDCA, and taurochenodeoxycholic acid; TCDCA) as well as 5-ASA were supplemented in the culture medium at UCSD (Zengler lab) and Cedars-Sinai labs to evaluate the production of bile acid–5-ASA conjugates. Bacterial cultures were incubated for 72 h in a shallow 96-well plate with edges filled with medium to prevent evaporation. Cultures were transferred to a 2 mL 96-well plate, extracted overnight at 4 ℃ for time = 0 h (baseline) and at time = 72 h (end of incubation) with 80% pre-chilled MeOH/H_2_O at a 4:1 ratio (solvent:sample). Samples were centrifugated at 2,500 RPM, then 600 µL was dried in a CentriVap overnight and stored at-80 ℃ until LC-MS/MS analysis. For the culturing done at MUSC, bacteria were grown in BHI broth overnight at 37 °C. After confirming growth, cultures were centrifuged at 6,000 x *g* for 5 min to pellet bacteria and the bacterial pellets were washed 3x with sterile PBS. The bacterial pellets were then resuspended in an equal volume of a chemically defined culture medium called ZMB1^80^ and sub-cultured to an optical density (OD_600nm_) of 0.1 in 5 mL of ZMB1 in triplicate with or without 0.01% bovine bile and 12.5 µg/mL 5-ASA. After 20 h of incubation, the cultures were centrifuged at 5,000 x *g* for 5 min to pellet the bacteria and the conditioned media supernatant samples were sterile filtered using 0.2 μm filters and processed for 5-ASA and cholyl–5-ASA bioanalysis. For the culturing of *Bifidobacterium* (UCSD Raffatellu/Chu labs), all isolates were grown on MRS agar plate and incubated overnight to check for pure cultures. *Bifidobacterium* isolates were inoculated into MRS broth supplemented with 1 mM cysteine, hemin, and vitamin K1 and incubated at 37 ℃ for 72 h. Cells were washed using fresh MRS broth and resuspended in MRS supplemented with 1 mM cysteine, hemin, and vitamin K1. Cells were normalized at an OD_600nm_ of 0.02 before transferring 200 µL of cells in a 96-well plate. Bile acids and 5-ASA were supplemented to the growth medium. After 72 h of growth, cultures were transferred to a 2 mL 96-well plate, extracted overnight at 4 ℃ for Time = 0 h (baseline) and at Time = 72 h (end of incubation) with 80% pre-chilled MeOH/H_2_O at a 4:1 ratio (solvent:sample). Samples were centrifugated at 975 x *g*, then 600 µL was dried in a CentriVap overnight and stored at-80 ℃ until LC-MS/MS analysis. The data is publicly available on MassIVE (MSV000100265).

### BSH screening assay and metabolite extraction

The *bsh*-expressing *Escherichia coli* were cultured overnight at 37 ℃ with shaking in 3 mL BHI. Overnight cultures were diluted 1:100 into BHI and grown for 4 h at 37 ℃ with shaking, after which cultures were normalized to an OD_600_ of 0.02. Control conditions included *E. coli* engineered to express green-fluorescent protein (GFP) and medium-only controls with substrates without bacteria. A 10 mM stock solution of TCA, TDCA, and TLCA were prepared in DMSO and diluted to a final concentration of 100 µM. Also, a 20 mM stock solution of 5-ASA was prepared in DMSO and diluted to 200 µM final concentration. A baseline (Time = 0 h) was established by immediately freezing the plate at-80 ℃. The cultures were grown at 37 ℃ in static condition for 48 h before freezing the plate at-80 ℃. Samples were thawed at room temperature for 30 min to 1 h. The whole culture (200 µL) was transferred to a 96-deep well plate and 800 µL of 100% MeOH containing 1 µM of sulfamethizole was added, followed by an overnight incubation at 4 ℃. Samples were centrifuged at 975 x *g* for 10 min at 4 ℃ and 600 µL was transferred and dried in a CentriVap overnight. Samples were resuspended in 50% MeOH/H_2_O containing 1 µM of sulfadimethoxine and incubated at-20 ℃ for 1 h. Samples were again centrifuged at 975 x *g* and 150 µL was transferred into a shallow 96-well plate for LC-MS/MS analysis. A list of the cloned BSH is available in Supplementary Table S5. Also, the metabolomics data is available on MassIVE (MSV000100335).

### Untargeted metabolomics data acquisition

All samples acquired for this study were randomized and run on an Orbitrap Exploris 240 coupled to a Vanquish UHPLC system (Thermo Fisher Scientific). Chromatographic separation was performed using a polar C18 (Kinetex, 100 x 2.1 mm, particle size 2.6 µm, pore size of 100 Å, Phenomenex). Five microliters of sample were injected and eluted at a flow rate of 0.5 mL/min using the following gradient: 0.5-1 min 5%B; 1-7.5 min 40% B; 7.5-8.5 min 99%B; 8.5-9.5 min 99%B; 9.5-10 min 5%B; 10-10.5 min 5%B; 10.5-10.75 min 99%B; 10.75-11.25 min 99% B; 11.25-11.5 min 5% B; 11.5-12 min 5%B. MS/MS spectra were acquired in data-dependent acquisition (DDA) mode using positive ionization. ESI+ parameters were set to 3.5 kV spray voltage, sheath gas to 50 Arb, auxiliary gas to 10 Arb, sweep gas to 1 Arb, ion transfer tube 325℃, and 350 ℃ for vaporizer temperature. The mass scan range was set to 150-1500 *m/z* with a resolution of 60,000 at *m/z* 200 with one microscan. RF lens 70%, AGC target set to standard, maximum injection time set to auto, and the data acquired in profile mode. An intensity of 1E4 was set as the threshold to trigger a DDA scan. Dynamic exclusion was set to custom. Precursor was excluded after two times if it occurs within 3 s. Exclusion duration was set to 5 s. For DDA MS2 scan, the isolation window was set to 1 *m/z* and the isolation offset to OFF. The normalized collision energies were set to a stepwise increase of 25, 40, and 60%. Top N was set to 10 per MS1 scan acquired with a resolution at *m/z* 200 of 22,500 with one microscan. Scan range mode set to auto, AGC target set to custom with 200% as normalized AGC target, maximum injection time set to auto, and the spectra was acquired in profile mode.

### LC-MS/MS data processing

The raw files generated by the mass spectrometers were converted to an open-access format mzML using MSConvert (version 3.0.25042 64-bit). Data generated in this study have been deposited in GNPS/MassIVE under the accession number MSV000097269. Feature extraction and detection were performed using the batch mode in MZmine 4. The batch files used to process the mzML files can be found in GitHub (data availability section). The advanced import option was used for mass detection and the factor of the lowest signal was used with a noise factor of 3 for MS1 and 2 for MS2. Chromatogram builder parameters were set to a minimum of 5 consecutive scans, minimum intensity and absolute height of 1E4 with a *m/z* tolerance of 10 ppm. Smoothing was applied using the Savitzky Golay algorithm using a retention time width of 5 s. Local minimum resolver parameters were set to a chromatographic threshold of 90%, minimum scan range of 0.05, a minimum absolute height of 1E4, peak top/edge ratio of 1.2, and a minimum scan of 3. 13C isotope filter and isotope peak finder were applied with a retention time tolerance of 0.04 min and a *m/z* tolerance of 3 ppm, respectively. Features were aligned using the join aligner module with a *m/z* tolerance of 10 ppm, weight *m/z* of 3, retention tolerance of 0.05 min. Feature list rows filter was applied with a minimum aligned features of 2 or 10% of the samples. Peak finder was set to an intensity tolerance of 20%, a *m/z* tolerance of 10 ppm, a retention time tolerance of 0.05 min, and a minimum of 3 scans. A duplicate peak filter was applied. metaCorrelate feature grouping was set with the following parameters: retention time tolerance of 0.06 min, minimum samples of 3, and a minimum Pearson correlation of 70% for feature height. Ion identity networking was also performed before exporting the final feature table. The GNPS and SIRIUS^84^ export module were used to create the csv file containing peak areas and the mgf files (MS/MS spectra list) necessary for downstream analysis in GNPS2 and R studio.

The output files from MZmine were used as input in the feature-based molecular networking workflow available in GNPS2. The parameters used were the following: precursor and fragment ion tolerance of 0.02, without filters, and with networking and library search set to cosine of 0.7 with a minimum of 6 matched peaks. The FBMN job is available can be accessed at:

Monoculture of gut microbes with 5-aminosalicylic acid and bile acids (MSV000097269): https://gnps2.org/status?task=ec12e706a21d4d178983c9380174fb82

### Data analysis of bacterial monocultures

Tables generated using MZmine were imported into R studio (version 4.5.0). Quality control of the run was assessed by inspecting the peak area and retention time drift as well as the coefficient of variance of the internal standard (sulfadimethoxine) present in the biological samples and a mixture of six different standards (QCmix; amitriptyline, sulfamethazine, coumarin 314, sulfadimethoxine, sulfamethizole, and sulfachloropyridazine) included every 10 samples during data acquisition using the table generated by the legacy MZmine 2 module. Features were removed from samples if they were not at least 5 times higher than the ones observed in pools and QCmix. Filtering steps were performed for the features of interest. A feature has to be detected in at least two biological replicates and for the radial network, known and unknown bile acids need to be at least 3 times higher at 72 h (after incubation) compared to time 0 h (baseline) to be visualized. The scripts for the analysis and to generate the figures can be found on GitHub https://github.com/VCLamoureux/5ASA_manuscript.

### Multi-omics data analysis

Metagenomics data acquired from the IBD200 cohort (Qiita #12675) was reprocessed. Detailed information on sample preparation and sequencing protocol can be found in the original publication^36^. Demultiplexed per-sample FASTQ files underwent a recently developed host filtration pipeline to remove human reads^52^. Briefly, reads mapping to reference human genomes (GRCh38.p14 + Phi X 174 + T2T-CHM13v2.0) via minimap were discarded, then the reads were subsequently filtered using a pangenome index generated from 94 reference assemblies from the Human Pangenome Reference Consortium^85^ using Movi^86^. Cleaned FASTQ files were then processed via the default shotgun metagenomics processing pipeline in Qiita^87^, which consists of applying bowtie2^89^ followed by woltka to align reads against the Web of Life (WoL2)^90^ reference database. Then, reads were filtered against Greengenes2 (v. 2024.09)^91^ to generate the final OGU table. Additionally quality filters were then applied. OGUs with <5% genome coverage, calculated via micov^92^, and present <5% of the samples were discarded. Additionally, samples with less than 50,000 cumulative reads were removed from downstream analysis. Average reads per sample was 960,709. Matching untargeted metabolomics data acquired from the IBD200 cohort (MSV000084775) was also reprocessed via MZmine as previously described. After blank subtraction, features detected in <5% of the samples were removed. Data was then subset to retain only samples in which 5-ASA was detected and for which matching metagenomics data was available (n=53). Multi-omics analysis was then performed on the filtered matching microbiome and metabolomics tables via joint-RPCA^53^. The obtained correlation matrix was then subsetted to extract OGUs positively correlating with cholyl-5-ASA. Correlation values were then collapsed to the mean by genera and the total number of OGUs of interest per genera was calculated to be visualized via a balloon plot.

### Animals

All animal studies were approved by the Institutional Animal Care and Use Committee at the Medical University of South Carolina (MUSC). Adult 12-16 week old male C57BL/6J mice (n = 4-5/group) were administered 3% DSS in their drinking water ad libitum. As a control, one set of mice received normal drinking water. The mice receiving DSS were randomly assigned into three groups: (1) 0.5% carboxymethylcellulose sodium in water (vehicle control), (2) 75 mg/kg of 5-ASA dissolved in 0.5% carboxymethylcellulose sodium or (3) 75 mg/kg of cholyl–5ASA dissolved in 0.5% carboxymethylcellulose sodium. Mice were monitored daily for weight loss and euthanized on day 5. At the time of collection, the ileum and colon were excised and flushed with 0.5 mL of sterile PBS for luminal 5-ASA and cholyl–5ASA bioanalysis. The median lobe of the liver was also collected for 5-ASA and cholyl–5ASA bioanalysis. The flushed colon was fixed in 10% neutral-buffered formalin at 4 °C overnight and paraffin-embedded. Tissue sections were stained H&E to examine colonic architecture and histological scores of colitis were assessed by a blinded pathologist.

### PPAR-γ activity assay

To test whether 5-ASA and cholyl–5-ASA can induce PPAR-γ activation, we used a reporter assay in proprietary rodent cell line that is engineered to express the Mouse / Rat Peroxisome Proliferator-Activated Receptor Gamma (nr1c3, pparG). This assay is designed to measure the transactivation potential of nuclear receptors independently of tissue-specific physiological context. The mousePPAR-γ reporter assay kit from Indigo Biosciences (mPPARγ, MR0010) was used to assess the effect of 5-ASA and cholyl–5-ASA. Assays were performed according to the manufacturer’s instructions. Briefly, the PPAR-γ reporter cell suspension was thawed by adding the cell recovery medium (CRM), and 100 µL of cell suspension were dispensed per well, followed by the addition of 100 µL of compound screening medium supplemented with test compounds. Cells were incubated at 37 ℃ with 5% CO_2_ for 24 h. PPAR-γ luminescence was quantified using the GloMax® luminometer (Promega) with Instinct software version 3.1.3. Docosahexaenoic acid (DHA) was assayed from 100 µM to 0.78 µM and rosiglitazone from 1 µM to 0.5 nM as a positive controls; 5-ASA, cholyl–5-ASA, and cholic acid were assayed from 62.5 µM to 2.5 mM. Cell toxicity and viability were assessed using the MTT cell proliferation assay kit (Abcam; Ab-211091).

### Immunomodulatory activity

Co-culture of bone marrow-derived dendritic cells (BMDCs) and bulk CD4^+^ T cells were performed previously described. Briefly, BMDCs were generated from bone marrow progenitor cells isolated from femurs of WT C57BL6/J mice in the presence of 20 ng/mL GM-CSF (Miltenyi) in complete RPMI 1640 (10% fetal bovine serum, 50 U/mL penicillin, 50 µg/mL streptomycin, 2 mM L-glutamine, 1 mM sodium pyruvate, 1 mM HEPES, non-essential amino acids, and beta-mercaptoethanol). BMDCs co-cultured with splenic CD4^+^ T cells at a ratio of 1:10 (DC:CD4^+^ T cells) and treated with 50 and 100 µM of bile acids in the presence of anti-mouse CD3 (2 μg/mL, eBiosciences), mouse IL-2 (10 μg/mL, Peprotech), mouse IL-6 (20 ng/mL, Peprotech), mouse IL-1b (20 ng/mL, Peprotech), mouse IL-23 (10 ng/mL, Life Technologies) and human TGF-beta (2 ng/mL, Peprotech). After three days of co-culture, cells were stimulated with a cell activation cocktail with Brefeldin A (2 µL/mL, Biolegend) for 4 h and stained for 30 min at 4 °C with either LIVE/DEAD fixable yellow dead stain Kit (Life Technologies), with empirically titrated concentrations of the following antibodies: eFluor506 anti-mouse CD4 (clone: RM4-5). For intracellular staining, cells were fixed and permeabilized using the Foxp3/Transcription Factor Staining Buffer Set (Life Technologies) according to the manufacturer’s protocol. Intracellular staining was performed with BrilliantViolet785-conjugated anti-mouse IFNγ (clone: XMG1.2), and FITC anti-mouse Foxp3 (clone: FJK-16s) for 4 h. All antibodies were purchased from Thermo Scientific/eBiosciences, and Biolegend. Cell acquisition was performed on Fortessa (BD), and data was analyzed using FlowJo software suite (TreeStar).

### Chemicals, consumables, and supplies

Honeywell – Burdick and Jackson produced LC-MS grade water, methanol, and acetonitrile solvents and Optima™ LC/MS-grade formic acid were purchased from Fisher Scientific (Waltham, MA, USA). Dimethyl sulfoxide (DMSO) solvent, LiChropur™ ammonium formate, and dextran sodium sulfate (DSS), and carboxymethylcellulose sodium were purchased from Millipore-Sigma (Burlington, MA, USA). The 2 mL fast prep-tubes (Catalog #MP115076200) and Lysing Matrix D 1.4 mm ceramic beads (Catalog #116540434) were purchased from MP Biomedical (Irvine, CA, USA). The 5-ASA standard (Catalog #A229125G) and bovine bile (Catalog # 50493748) were purchased from Fisher Scientific. The cholyl–5-ASA standard was synthesized at University of California - San Diego. The analytical and guard column combination used for the cholyl–5-ASA method was a 2.7-micron Raptor C18 (End-Capped; 2.1 mm x 100 mm; 90 Å pore; Catalog # 9304A12) analytical column coupled to an 5-micron Ultra C18 (2.1 mm x 10 mm; Catalog # 917850212) guard column purchased from Restek (Bellefonte, PA, USA). The mixed-mode analytical column used for the 5-ASA method was a 3.0-micron Amaze MH (2.1 x 30 mm, 100Å pore; Catalog #AMHII1505) purchased from Helix Chromatography (Prospect Heights, IL, USA).

### Retention time matching

Retention time matching and MS/MS spectral similarities were performed between the biological sample and the crude synthetic standard available on MassIVE (MSV000100244). Fecal samples from the RA cohort (MSV000084556; IRB#161474) were available in our laboratory. Briefly, the sample (45.6 mg) was transferred into a pre-weighted Eppendorf tube containing a stainless steel 5 mm and extracted using pre-chilled 50% MeOH/H_2_O at a ratio of 50 mg of sample to 800 µL of solvent. The sample was homogenized in a Qiagen TissueLyzer II for 5 min at 25 Hz and incubated at 4 °C for 1 h before centrifugation at 15,000 x *g* for 10 min. The supernatant (200 µL) was dried overnight in a CentriVap and stored at-80 °C before resuspension. The cholyl–5-ASA and the lithocholyl–5-ASA crude combinatorial synthesis were injected on an Agilent system coupled to a hybrid trapped ion mobility-quadrupole time-of-flight mass spectrometer (timsTOF pro II). A kinetex (Phenomenex) non-polar C18 column (2.1 mm x 100 mm, 2.6 µm particle size 100 A pore size) was used with a guard cartridge (2.1 mm ID) matching the column type. The mobile phase consisted of solvent A (H_2_O + 0.1% formic acid) and the solvent B (ACN + 0.1% formic acid) and the column compartment was heated at 40 °C. Five microliters of samples were injected and eluted at a flow rate of 0.5 mL per min using the following gradient: 0 – 0.5 min 5% B, 0.5 – 1.1 min 25% B, 1.1 – 7.5 min 40% B, 7.5 – 8.5 min 99% B, 8.5 – 10 min 99% B, 10 – 10.1 min 5% B, 10.1 – 12 min 5% B. The QTOF was pre-calibrated using NA formate in positive mode at ppm standard deviation below 0.5%. Mass spectra were recorded at *m/z* 20-1300 and scan mode PASEF. End plate offset was set to 500 V, capillary to 4500 V, Nebulizer on at 2.2 bar, dry gas at 10 L/min, and dry temperature at 200 °C. The low-abundance precursor ions with an intensity above 100 counts but below a target value of 4000 counts were scheduled with an active exclusion of 0.05 min. Collision energies were set to 50 and 60 eV. The deoxycholyl–5-ASA was injected on a Vanquish UHPLC (ultra-high performance liquid chromatography) system coupled to a Q-Exactive Orbitrap mass spectrometry (Thermo Fisher Scientific). The mobile phase A and B were H_2_O + 0.1% formic acid and ACN + 0.1% formic acid, respectively. A kinetex (Phenomenex) non-polar C18 column (2.1 mm x 100 mm, 2.6 µm particle size 100 A pore size) was used with a guard cartridge (2.1 mm ID) matching the column type and the column compartment was heated at 40 °C. Five microliters were injected and eluted using the following gradient: 0-1 min 5 % B, 1-7 min 5-40 % B, 7-10 min 40-98% B, and 10-12 min 5% B for re-equilibration. A positive mode electrospray ionization was used with the following parameters: sheath gas flow, 53 AU; auxiliary gas flow, 14 AU; sweep gas flow, 3 AU; auxiliary gas temperature, 400 °C; spray voltage, 3.5 kV; inlet capillary temperature, 269 °C; S-lens level, 50 V. MS1 scan was performed at *m/z* 100-1500 with the following parameters: resolution, 17,500 at *m/z* 200; maximum ion injection time, 100 ms; automatic gain control (AGC) target, 1E6. Up to 5 MS/MS spectra per MS1 scan were recorded under the data-dependent acquisition mode (DDA) with the following parameters: resolution, 17,500 at *m/z* 200; maximum ion injection time, 150 ms; AGC target, 5E5; isolation window, *m/z* 1; isolation offset, *m/z* 0.5; stepped normalized collision energy (NCE) of 25%, 40%, and 60%; minimum AGC target 5E4; apex trigger, 2 to 5 s; dynamic precursor exclusion set to 10 s.

### Instruments for quantitation of 5-ASA and cholyl-5-ASA

The LC-HRMS system is comprised of a Nexera Series-40 ultrahigh-performance liquid chromatography (UHPLC) system outfitted with a SIL-30ACMP autosampler (Shimadzu, Kyoto, Japan) that is coupled to a ZenoTOF 7600+ quadrupole-time-of-flight MS (QTOF MS) system (SCIEX, Framingham, MA, USA). The system was operated using SCIEX OS software (Version 3.3.1.43), while peak integration and quantitative analysis were performed using the Analytics module in SCIEX OS.

### 5-aminosalicylic acid stock, intermediate, infusion, and calibrator preparations

A 5-ASA stock solution was prepared at 10 mg/mL in pure DMSO and was used to prepare an intermediate solution that contained 100 µg/mL of 5-ASA in a solution consisting of methanol: water (1:1, *v:v*). This intermediate solution was used to prepare an infusion solution that contained 100 ng/mL of 5-ASA in a solution consisting of methanol: water (1:1, *v:v*); this solution was infused at a rate of 10 µL/min into the ionization source of the ZenoTOF 7600+ to identify the 5-ASA-based [M+H]+ precursor ion, all diagnostic 5-ASA product ions, and to optimize all of the 5-ASA-specific parameters for the ZenoTOF 7600+ (See the section titled LC-HRMS method for 5-ASA quantitative analysis and **Supplementary Table S6**).

The intermediate solution was also used to prepare a high calibrator that contained 10,000 ng/mL of 5-ASA in a solution consisting of acetonitrile: water (9:1, *v:v*). A 4-fold serial dilution procedure was used to produce an 8-point calibration curve composed of calibrators that contained 5-ASA concentrations of 0.610 ng/mL, 2.44 ng/mL, 9.76 ng/mL, 39.06 ng/mL, 156 ng/mL, 625 ng/mL, 2,500 ng/mL, and 10,000 ng/mL per level using a solution consisting of acetonitrile:water (9:1, v:v) as the diluent.

### 5-ASA mobile phase and needlewash preparations

5-ASA separations were performed using a mixed-mode chromatographic approach using an Amaze MH analytical column. Mobile phase A (MPA), mobile phase B (MPB) and needlewash (NW) preparations consisted of a mixture of acetonitrile: 100 mM ammonium formate (pH 4) solution (9:1, *v:v*), a mixture of acetonitrile: 100 mM ammonium formate (pH 4) solution: 3% formic acid solution (50/5/45, *v:v:v*), and a mixture of acetonitrile: water (1:1, *v:v*), respectively. After thorough mixing, the MPA, MPB and NW solutions were each degassed using pulsed bath sonication (Elmasonic S, Elma Schmidbauer GmbH, Singen, Germany) for 15 min without heating prior to use. These mobile phase solutions were stored well sealed at ambient temperature and expired two months after preparation.

### LC-HRMS method for 5-ASA quantitative analysis

The UHPLC system was specified to deliver a mobile phase flowrate of 0.600 mL/min at an initial mobile phase composition of 95:5 MPA: MPB, the analytical column was heated to 40 °C, and the mobile phase gradient was programmed as follows: 0.00-0.25 min, 5% MPB; 0.25-2.00 min, 5-50% MPB; 2.00-2.25 min, 50% MPB; 2.25-2.30 min, 50-5% MPB; 2.30-4.00 min, 5% MPB with a per sample cycle time of ∼4.4 min. The SIL-30ACMP autosampler settings were specified as follows: Sampling speed, 5.0 µL/s; Cooler temperature, 8°C; Temperature control, enabled for Rack plate L, Rack plate M, and Rack plate R; Rinse type, External only; Rinse mode, Before and after aspiration; Dip time: 0 s; Rinse method, Rinse port only; Time, 2 s; Sampling speed, 35 µL/s; Rinse volume, 500 µL; Measuring line purge volume, 100 µL. The retention time (RT) for 5-ASA was 1 min using these conditions.

The ZenoTOF 7600+ was outfitted with a ZenoTOF TwinSpray probe in a TurboV Source, and the global mass spec parameters were specified as follows: Workflow, Small Molecule; Method duration, 4 min; Total scan time, 0.266 s (software determined); and, Estimated cycles, 901 (software determined). The Source and Gas Parameters were specified as: Ion source gas 1, 50 psi; Ion source gas 2, 70 psi; Curtain gas, 40 psi; CAD gas, 9; Temperature, 500 °C. The Experiment scan was set to MRMHR; Polarity, Positive; Spray voltage, +5,500 V; TOF start mass, 50 Da; TOF stop mass, 1000 Da; Accumulation time, 0.030 s; Declustering potential, 30 V; DP spread, 0 V; Collision Energy, 12 eV; CE spread, 0 eV. Advanced Experimental Settings included a Time bins to sum setting of 6, Channels 1-4 were enabled, and the system selected QJet amplitude was 183.248 V. The TOF MSMS settings included: Q1 resolution, Unit; Zeno Pulsing, enabled; Zeno threshold, 150000 CPS; Manual QJet RF amplitude, disabled; QJet RF amplitude, 94.972 V. The 5-ASA-specific MRMHR-based TOF MSMS parameters were specified as follows: [M+H]+ precursor ion, *m/z* 154.04; TOF Start Mass (Da), *m/z* 20.0000; TOF Stop Mass (Da) *m/z* 150.0000; Accumulation Time, 0.0400 s; Declustering Potential, +30 V; CE Spread, 0 eV; Time bins to sum, 6. The Collision Energies are tabulated in Supplementary Table S6.

### Cholyl–5-ASA stock, intermediate, infusion, and calibrator preparations

A cholyl–5-ASA stock solution was prepared at a concentration of 2.5 mg/mL in a solution consisting of methanol: water (1:1, *v:v*) and was used to prepare an intermediate solution that contained 5 µg/mL of cholyl-5-ASA in a solution consisting of methanol: water (1:1, *v:v*). This intermediate solution was used to prepare an infusion solution that contained 500 ng/mL of cholyl-5-ASA in a solution consisting of methanol: water (1:1, *v:v*); this solution was infused at a rate of 20 µL/min into the ionization source of the ZenoTOF 7600+ to identify the cholyl-5-ASA-based [M-H]-precursor ion, all diagnostic cholyl–5-ASA product ions, and to optimize all of the cholyl–5-ASA specific parameters for the ZenoTOF 7600+ (See the section titled LC-HRMS method for cholyl–5-ASA quantitative analysis and Supplementary Table S7).

The intermediate solution was also used to prepare a high Calibrator that contained 500 ng/mL of cholyl–5-ASA in a solution consisting of methanol: water (1:1, *v:v*). A 4-fold serial dilution procedure was used to produce a 6-point calibration curve composed of calibrators that each contained cholyl–5-ASA concentrations of 0.488 ng/mL, 1.95 ng/mL, 7.81 ng/mL, 31.25 ng/mL, 125 ng/mL, and 500 ng/mL per level using a solution consisting of methanol: water (1:1, *v:v*) as the diluent.

### Cholyl–5-ASA mobile phase and needlewash preparations

Cholyl–5-ASA separations were performed using a reverse phase chromatographic approach using a Raptor C18 analytical and Ultra C18 guard column combination. MPA, MPB and NW solutions consisted of 10 mM ammonium formate (unbuffered) solution, pure acetonitrile and pure isopropanol, respectively. These mobile phase solutions were stored well sealed at ambient temperature and expired two months after preparation.

### LC-HRMS method for cholyl–5-ASA quantitative analysis

UHPLC system parameters included a flowrate of 0.3 mL/min at an initial mobile phase proportion of 75:25 MPA: MPB, column temperature of 50 °C, and the mobile phase gradient was programmed as follows: 0.0-1.5 min, 25% MPB; 1.5-13.0 min, 25-80% MPB; 13.0-14.5 min, 80% MPB; 14.5-14.6 min, 80-25% MPB; 14.6-17.0 min, 25% MPB with a per sample cycle time of ∼17.4 min.

The SIL-30ACMP autosampler settings were specified as follows: Sampling speed, 5.0 µL/s; Cooler temperature, 10 °C; Temperature control, enabled for Rack plate L, Rack plate M, and Rack plate R; Rinse type, External only; Rinse mode, Before and after aspiration; Dip time: 0 s; Rinse method, Rinse port only; Time, 2 s; Sampling speed, 35 µL/s; Rinse volume, 500 µL; and, Measuring line purge volume, 100 µL. The retention time (RT) for cholyl–5-ASA was 5.15 min using these conditions.

The ZenoTOF 7600+ was outfitted with a ZenoTOF TwinSpray probe in a TurboV Source, and the global mass spec parameters were specified as follows: Workflow, Small Molecule; Method duration, 17 min; Total scan time, 0.215 s (software determined); and, Estimated cycles, 4735 (software determined). The Source and Gas Parameters were specified as: Ion source gas 1, 50 psi; Ion source gas 2, 70 psi; Curtain gas, 40 psi; CAD gas, 9; Temperature, 500 °C. The Experiment scan was set to MRMHR; Polarity, Negative; Spray voltage,-4,500 V; TOF start mass, 80 Da; TOF stop mass, 1000 Da; Accumulation time, 0.100 s; Declustering potential,-60 V; DP spread, 0 V; Collision Energy,-12 eV; and, CE spread, 0 eV. Advanced Experimental Settings included a Time bins to sum setting of 6, Channels 1-4 were enabled, and the system selected QJet amplitude was 183.246 V. The TOF MSMS settings included: Q1 resolution, Unit; Zeno Pulsing, enabled; Zeno threshold, 20000 CPS; Manual QJet RF amplitude, disabled; and the system selected QJet RF amplitude, 202.72 V. The cholyl–5-ASA specific MRMHR-based TOF MSMS parameters were specified as follows: [M-H]-precursor ion, *m/z* 542.31; TOF Start Mass (Da), *m/z* 80.0000; TOF Stop Mass (Da) *m/z* 1000.0000; Accumulation Time, 0.0100 s; Declustering Potential,-60 V; CE Spread, 0 eV; Time bins to sum, 6. The precursor ion, diagnostic product ions, and collision energies are tabulated in Supplementary Table S7.

### Synthesis

NMR spectra were collected at 298 K on a 600 MHz Bruker Avance III spectrometer fitted with a 1.7 mm triple resonance cryoprobe with z-axis gradient (^1^H NMR: MeOD (3.31), CDCl_3_ (7.26) at 500 MHz). Cholyl–5-ASA spectra was collected in MeOD with shifts reported in parts per million (ppm) referenced to the proton of the solvent (3.31). Data for ^1^H-NMR are reported as follows: chemical shift (ppm, reference to protium; s = single, d = doublet, t = triplet, q = quartet, dd = doublet of doublets, m = multiplet, coupling constant (Hz), and integration). **Cholyl–5-ASA** ^1^H NMR (600 MHz, MeOD) δ 7.65 (s, 1H), 7.47 (d, 1H), 6.65 (d, 1H), 3.70 (d, 1H), 3.87 (s, 1H), 3.70 (d, 1H), 2.27-2.31 (m, 3H), 1.76-1.93 (m, 5H), 1.60 – 1.71 (m, 2H), 1.39 – 1.55 (m, 6H), 1.24 – 1.138 (m, 5H), 1.97 (m, 3H), 1.65 (m 1H), 1.63-1.50 (m, 3H), 1.47 – 1.33 (m, 1H), 0.85 – 1.05 (m, 1H), 0.82 (d, 3H), 0.99 (m, 1H), 0.82 (s, 3H), 0.65 (s, 3H). The NMR spectrum is available in **Supplementary Figure S5.**

### Cholyl–5-ASA synthesis

Solid cholic acid (2 mmol, 1 g, 1 eq.) and 5 mL of THF were added to a 20 mL glass vial with a stir bar and the reaction vessel was placed in an ice water bath. Neat ethyl-chloroformate (3 mmol, 300 µL, 1.2 eq.) and triethylamine (4 mmol, 500 μL, 1.5 eq.) were subsequently added, and the solution was stirred for 1.5 h at 0 °C. 5-aminosalicylic acid (2 mmol, 400 mg, 1 eq.) was dissolved in 5 mL of 1 M NaOH and subsequently added in one portion to the cholic acid mixture. The reaction was stirred overnight at 23 °C. The mixture was then concentrated *in vacuo* and purified by CombiFlash NextGen 300+ using reversed phase column C18 15.5 g Gold at a flow rate of 13 mL per min with H_2_O (Solvent A) and ACN (solvent B) using the following gradient: 0-5 min, 5% B; 5-14 min, 40% B; 14-20 min 40% B; 20-25 min, 80% B. Cholyl–5-ASA eluted at 15 min, 40% B.

## Data availability

Untargeted metabolomics datasets reanalyzed in this study are publicly available in GNPS/MassIVE and includes: MSV000084556 (rheumatoid arthritis cohort), MSV000084775 (IBD cohort 1), MSV000082094 (IBD cohort 2), MSV000097610 (paediatric IBD cohort), and MSV000097269 (microbial culture with 5-ASA). iPENS (IRB#19/WS/0163) data, due to UK GDPR regulations, are available upon request to Konstantinos Gerasimidis and will require a data transfer agreement.

## Code availability

All the scripts used to perform data analysis and generate figures are publicly available at https://github.com/VCLamoureux/5ASA_manuscript.

## Acknowledgements

PCD acknowledges CCFA 1243263, NIDDK U24DK133658, NIAID R37AI126277, R21AR082029, R24GM148372, NIDDK R01DK136117, and Chan Zuckerberg Initiative for funding that all enabled parts of this work. V.C.L. is supported by Fonds de recherche du Québec - Santé (FRQS) postdoctoral fellowship (335368) and from Natural Sciences and Engineering Research Council of Canada (NSERC) postdoctoral fellowship (598938). The Texas Children’s Hospital Department of Pathology provides salary support to the bioanalytical chemistry staff members who work in The Virginia and L.E. Simmons Family Foundation Mass Spectrometry laboratory within the Texas Children’s Research Institute – Microbiome Center, and purchased all of reagents, consumables and durable supplies described in the targeted metabolomics sections of this manuscript. SFHS is supported by the National Institute of Health (F32DE035388-01). MHL was supported by the National Institute of Health (F32 AI169989). GTW was supported by NIH training grant T32AI007036. GJN was supported by NIH training grant T32 DK007202 and grant 1F31AI186410-01. SG is supported by a Medical University of South Carolina (MUSC) Odyssey Fellowship. MAE is supported by the National Institute of Health (P30DK123704, P20GM120457, R35GM155451). TDH is supported by an Office of Research Infrastructure Programs (ORIP)-based S10 Award (NIH S10OD036416), the Texas Medical Center - Digestive Disease Center (NIH P30DK056338), and receives salary support from an NIAID multicenter P01 grant (P01AI152999). LP is supported by the University of California - San Diego Medical Scientist Training Program (NIH/NIGMS T32GM007198). MG is supported by the Krupp Endowed Fund. JGM acknowledges funding from the NIGMS R35GM155339, MPRINT supported by NIH grant P30HD106451 and Vanderbilt University Medical Center’s Digestive Disease Research Center supported by NIH grant P30DK058404. RH lab is supported by The Archie Foundation and RH has research time supported by Tayside Medical Science Centre. ADP was supported by NIH/National Institute of Environmental Sciences grant R35 ES035027, and the US Department of Agriculture National Institute of Food and Federal Appropriations under project PEN047702 and accession number 7006412. ERR was supported by PSU/NIDDK (T32DK120509). MR was supported by R37AI126277. JV is supported by NIH funding (P30 DK056338). NJWR was supported by Royal Society Industry Fellowship IF\R1\231034. HNZ was supported by the National Institute Of Environmental Health Sciences of the National Institutes of Health under Award Number K99ES037746.

## References

1. Cong, J. et al. Bile acids modified by the intestinal microbiota promote colorectal cancer growth by suppressing CD8+ T cell effector functions. Immunity 57, 876–889.e11 (2024).

2. Fu, T., et al. Paired microbiome and metabolome analyses associate bile acid changes with colorectal cancer progression. Cell Reports 42, (2023).

3. Mulak, A. Bile Acids as Key Modulators of the Brain-Gut-Microbiota Axis in Alzheimer’s Disease. Journal of Alzheimer’s Disease 84, 461–477 (2021).

4. Chiang, J. Y. L., Ferrell, J. M., Wu, Y. & Boehme, S. Bile Acid and Cholesterol Metabolism in Atherosclerotic Cardiovascular Disease and Therapy. Cardiol Plus 5, 159–170 (2020).

5. Nie, Q. et al. Gut symbionts alleviate MASH through a secondary bile acid biosynthetic pathway. Cell 187, 2717–2734.e33 (2024).

6. Ma, H. & Patti, M. E. Bile acids, obesity, and the metabolic syndrome. Best Practice & Research Clinical Gastroenterology 28, 573–583 (2014).

7. Lamichhane, S. et al. Microbiome-derived bile acid signatures in early life and their association with islet autoimmunity. Nature Communications 10.1038/s41467-025-66619-6 (2025).

8. Ferrell, J. M. & Chiang, J. Y. L. Short-Term Circadian Disruption Impairs Bile Acid and Lipid Homeostasis in Mice. Cellular and Molecular Gastroenterology and Hepatology 1, 664–677 (2015).

9. Thomas, J. P., Modos, D., Rushbrook, S. M., Powell, N. & Korcsmaros, T. The Emerging Role of Bile Acids in the Pathogenesis of Inflammatory Bowel Disease. Frontiers in Immunology Volume 13–2022, (2022).

10. Li, T. & Apte, U. Chapter Nine - Bile Acid Metabolism and Signaling in Cholestasis, Inflammation, and Cancer. in Advances in Pharmacology (ed. Hardwick, J. P.) vol. 74 263–302 (Academic Press, 2015).

11. Quinn, R. A. et al. Global chemical effects of the microbiome include new bile-acid conjugations. Nature 579, 123–129 (2020).

12. Mohanty, I. et al. The underappreciated diversity of bile acid modifications. Cell 187, 1801–1818.e20 (2024).

13. Kvitne, K. E. et al. Environmental and maternal imprints on infant gut metabolic development. Cell Host & Microbe doi:10.1016/j.chom.2025.11.002.

14. Zuffa, S. et al. A Multi-Organ Murine Metabolomics Atlas Reveals Molecular Dysregulations in Alzheimer’s Disease. bioRxiv 2025.04.28.651123 (2025) doi:10.1101/2025.04.28.651123.

15. Gentry, E. C. et al. Reverse metabolomics for the discovery of chemical structures from humans. Nature 626, 419–426 (2024).

16. Lin, J. et al. A microbial amino-acid-conjugated bile acid, tryptophan-cholic acid, improves glucose homeostasis via the orphan receptor MRGPRE. Cell 188, 4530–4548.e25 (2025).

17. Ramos, S. F. et al. Metatranscriptomics Uncover Diurnal Functional Shifts in Bacterial Transgenes with Profound Metabolic Effects. Cell Host Microbe 33, 1057–1072 (2025).

18. Wang, M. et al. Sharing and community curation of mass spectrometry data with Global Natural Products Social Molecular Networking. Nat Biotechnol 34, 828–837 (2016).

19. Charron-Lamoureux, V. et al. A searchable metadata network graph for microbiome metabolomics. Preprint at 10.64898/2026.02.04.703849 (2026).

20. Xing, S. et al. Navigating the conjugated metabolome. Preprint at 10.64898/2026.02.06.704496 (2026).

21. Hofmann, A. F. & Hagey, L. R. Key discoveries in bile acid chemistry and biology and their clinical applications: history of the last eight decades. Journal of Lipid Research 55, 1553–1595 (2014).

22. Haug, K. et al. MetaboLights: a resource evolving in response to the needs of its scientific community. Nucleic Acids Research 48, D440–D444 (2020).

23. Sud, M. et al. Metabolomics Workbench: An international repository for metabolomics data and metadata, metabolite standards, protocols, tutorials and training, and analysis tools. Nucleic Acids Research 44, D463–D470 (2016).

24. Batsoyol, N., Pullman, B., Wang, M., Bandeira, N. & Swanson, S. P-massive: a real-time search engine for a multi-terabyte mass spectrometry database. in Proceedings of the International Conference on High Performance Computing, Networking, Storage and Analysis 1–15 (IEEE Press, Dallas, Texas, 2022).

25. Wang, M. et al. Mass spectrometry searches using MASST. Nat Biotechnol 38, 23–26 (2020).

26. El Abiead, Y., et al. Enabling pan-repository reanalysis for big data science of public metabolomics data. Nature Communications 16, 4838 (2025).

27. Charron-Lamoureux, V. et al. A guide to reverse metabolomics—a framework for big data discovery strategy. Nature Protocols 10.1038/s41596-024-01136-2 (2025).

28. Lee, R. T. et al. Binding characteristics of N-acetylglucosamine-specific lectin of the isolated chicken hepatocytes: similarities to mammalian hepatic galactose/N-acetylgalactosamine-specific lectin. Biochemistry 28, 8351–8358 (1989).

29. Coras, R. et al. Baseline microbiome and metabolome are associated with response to ITIS diet in an exploratory trial in patients with rheumatoid arthritis. Clinical & Translational Med 12, e959 (2022).

30. Amos, R. S. The History of the use of Sulphasalazine in Rheumatology. British Journal of Rheumatology XXXIV, 2–6 (1995).

31. Klotz, U. Clinical Pharmacokinetics of Sulphasalazine, Its Metabolites and Other Prodrugs of 5-Aminosalicylic Acid. Clinical Pharmacokinetics 10, 285–302 (1985).

32. Pullar, T., Hunter, J. A. & Capell, H. A. Which component of sulphasalazine is active in rheumatoid arthritis? Br Med J (Clin Res Ed) 290, 1535–1538 (1985).

33. Khan, A. K. A., Piris, J. & Truelove, S. C. AN EXPERIMENT TO DETERMINE THE ACTIVE THERAPEUTIC MOIETY OF SULPHASALAZINE. The Lancet 310, 892–895 (1977).

34. Wörth, W. D. & Müller, W. [Effect of 5-aminosalicylic acid as a constituent of sulfasalazine in the treatment of chronic polyarthritis]. Z Rheumatol 45, 79–82 (1986).

35. Smedegård, G. & Björk, J. Sulphasalazine: Mechanism of Action in Rheumatoid Arthritis. British Journal of Rheumatology XXXIV, 7–15 (1995).

36. Mills, R. H. et al. Multi-omics analyses of the ulcerative colitis gut microbiome link Bacteroides vulgatus proteases with disease severity. Nature Microbiology 7, 262–276 (2022).

37. Zhao, H. N. et al. A resource to empirically establish drug exposure records directly from untargeted metabolomics data. Nat Commun 16, 10600 (2025).

38. Patan, A. et al. Charting the Undiscovered Metabolome with Synthetic Multiplexing. bioRxiv 2025.11.18.689170 (2025) doi:10.1101/2025.11.18.689170.

39. Kvitne, K. E. et al. Fecal microbial and metabolic signatures in children with very early onset inflammatory bowel disease. npj Biofilms Microbiomes (2025) doi:10.1038/s41522-025-00899-0.

40. Fang, X. et al. Gastrointestinal Surgery for Inflammatory Bowel Disease Persistently Lowers Microbiome and Metabolome Diversity. Inflammatory Bowel Diseases 27, 603–616 (2021).

41. Svolos, V. et al. European Crohn’s and Colitis Organisation consensus on dietary management of inflammatory bowel disease. Journal of Crohn’s and Colitis 19, jjaf122 (2025).

42. Svolos, V. et al. Treatment of Active Crohn’s Disease With an Ordinary Food-based Diet That Replicates Exclusive Enteral Nutrition. Gastroenterology 156, 1354–1367.e6 (2019).

43. Ventin-Holmberg Rebecka et al. The total gut mucosal and fecal bacterial load increases in successful treatment of inflammatory bowel disease with infliximab. Microbiology Spectrum 13, e01894–24 (2025).

44. Zheng, J. et al. Noninvasive, microbiome-based diagnosis of inflammatory bowel disease. Nat Med 30, 3555–3567 (2024).

45. Elmassry, M. M. et al. A meta-analysis of the gut microbiome in inflammatory bowel disease patients identifies disease-associated small molecules. Cell Host & Microbe 33, 218–234.e12 (2025).

46. Shan, Y., Lee, M. & Chang, E. B. The Gut Microbiome and Inflammatory Bowel Diseases. Annu Rev Med 73, 455–468 (2022).

47. Iliev, I. D., Ananthakrishnan, A. N. & Guo, C.-J. Microbiota in inflammatory bowel disease: mechanisms of disease and therapeutic opportunities. Nat Rev Microbiol 23, 509–524 (2025).

48. Becker, H. E. F., Demers, K., Derijks, L. J. J., Jonkers, D. M. A. E. & Penders, J. Current evidence and clinical relevance of drug-microbiota interactions in inflammatory bowel disease. Front. Microbiol. 14, 1107976 (2023).

49. Savage, H. P. et al. Epithelial hypoxia maintains colonization resistance against Candida albicans. Cell Host & Microbe 32, 1103–1113.e6 (2024).

50. Mehta, R. S. et al. Gut microbial metabolism of 5-ASA diminishes its clinical efficacy in inflammatory bowel disease. Nature Medicine 29, 700–709 (2023).

51. Weng, Y. et al. Calculating fast differential genome coverages among metagenomic sources using micov. Communications Biology 8, 1624 (2025).

52. Guccione, C. et al. Incomplete human reference genomes can drive false sex biases and expose patient-identifying information in metagenomic data. Nature Communications 16, 825 (2025).

53. Burcham, Z. M. et al. A conserved interdomain microbial network underpins cadaver decomposition despite environmental variables. Nature Microbiology 9, 595–613 (2024).

54. Rimal, B. et al. Bile salt hydrolase catalyses formation of amine-conjugated bile acids. Nature 626, 859–863 (2024).

55. Guzior, D. V. et al. Bile salt hydrolase acyltransferase activity expands bile acid diversity. Nature 626, 852–858 (2024).

56. Lucas, L. N. et al. Investigation of Bile Salt Hydrolase Activity in Human Gut Bacteria Reveals Production of Conjugated Secondary Bile Acids. Preprint at 10.1101/2025.01.16.633392 (2025).

57. Han, L. et al. Chemoproteomic profiling of substrate specificity in gut microbiota-associated bile salt hydrolases. Cell Chemical Biology 32, 145–156.e9 (2025).

58. Russell, B. J. et al. Intestinal transgene delivery with native E. coli chassis allows persistent physiological changes. Cell 185, 3263–3277.e15 (2022).

59. Cevallos, S. A. et al. 5-Aminosalicylic Acid Ameliorates Colitis and Checks Dysbiotic Escherichia coli Expansion by Activating PPAR-γ Signaling in the Intestinal Epithelium. mBio 12, e03227–20 (2021).

60. Rousseaux, C. et al. Intestinal antiinflammatory effect of 5-aminosalicylic acid is dependent on peroxisome proliferator–activated receptor-γ. The Journal of Experimental Medicine 201, 1205–1215 (2005).

61. Oh-oka, K. et al. Induction of Colonic Regulatory T Cells by Mesalamine by Activating the Aryl Hydrocarbon Receptor. Cellular and Molecular Gastroenterology and Hepatology 4, 135–151 (2017).

62. Di Paolo, M. C., Merrett, M. N., Crotty, B. & Jewell, D. P. 5-Aminosalicylic acid inhibits the impaired epithelial barrier function induced by gamma interferon. Gut 38, 115–119 (1996).

63. Britto, S. L., Krishna, M. & Kellermayer, R. Weight loss is a sufficient and economical single outcome measure of murine dextran sulfate sodium colitis. FASEB BioAdvances 1, 493–497 (2019).

64. Engevik, M. A., et al. *Bifidobacterium dentium*-derived y-glutamylcysteine suppresses ER-mediated goblet cell stress and reduces TNBS-driven colonic inflammation. Gut Microbes 13, 1902717 (2021).

65. Engevik, M. A. et al. Immunomodulation of dendritic cells by *Lactobacillus reuteri* surface components and metabolites. Physiol Rep 9, (2021).

66. Jordan, A. A. et al. Healthcare Access for Patients With Inflammatory Bowel Disease in the United States: A Survey by the Crohn’s & Colitis Foundation. Inflammatory Bowel Diseases 31, 1819–1832 (2025).

67. Updates on conventional therapies for inflammatory bowel diseases: 5-aminosalicylates, corticosteroids, immunomodulators, and anti-TNF-α FAU - Park, Jihye FAU - Cheon, Jae Hee. Korean J Intern Med 37, 895–905 (2022).

68. van de Meeberg, M. M., Schultheiss, J. P. D., Oldenburg, B., Fidder, H. H. & Huitema, A. D. R. Does the 5-Aminosalicylate Concentration Correlate with the Efficacy of Oral 5-Aminosalicylate and Predict Response in Patients with Inflammatory Bowel Disease? A Systematic Review. Digestion 101, 245–261 (2020).

69. Rousseaux, C. et al. Intestinal antiinflammatory effect of 5-aminosalicylic acid is dependent on peroxisome proliferator–activated receptor-γ. Journal of Experimental Medicine 201, 1205–1215 (2005).

70. Punchard, N. A., Greenfield, S. M. & Thompson, R. P. H. Mechanism of action of 5-arninosalicylic acid. Mediators of Inflammation 1, 480976 (1992).

71. Huang, L. et al. 5-Aminosalicylic acid ameliorates dextran sulfate sodium-induced colitis in mice by modulating gut microbiota and bile acid metabolism. Cellular and Molecular Life Sciences 79, 460 (2022).

72. Wang, J. et al. 5-aminosalicylic acid alleviates colitis and protects intestinal barrier function by modulating gut microbiota in mice. Naunyn-Schmiedeberg’s Arch Pharmacol 398, 3681–3695 (2025).

73. Tian, C. et al. The Efficacy and Safety of Mesalamine and Probiotics in Mild-to-Moderate Ulcerative Colitis: A Systematic Review and Meta-Analysis. Evidence-Based Complementary and Alternative Medicine 2020, 6923609 (2020).

74. Bahrami, S., Babaei, N., Esmaeili Gouvarchin Ghaleh, H., Mohajeri Borazjani, J. & Farzanehpour, M. Investigating the effects of combined treatment of mesalazine with Lactobacillus casei in the experimental model of ulcerative colitis. Front. Mol. Biosci. 11, 1456053 (2024).

75. Pardi, D. S., Loftus, E. V., Kremers, W. K., Keach, J. & Lindor, K. D. Ursodeoxycholic acid as a chemopreventive agent in patients with ulcerative colitis and primary sclerosing cholangitis. Gastroenterology 124, 889–893 (2003).

76. Batta, A. K., Tint, G. S., Xu, G., Shefer, S. & Salen, G. Synthesis and intestinal metabolism of ursodeoxycholic acid conjugate with an antiinflammatory agent, 5-aminosalicylic acid. Journal of Lipid Research 39, 1641–1646 (1998).

77. Goto, M., Okamoto, Y., Yamamoto, M. & Aki, H. Anti-inflammatory effects of 5-aminosalicylic acid conjugates with chenodeoxycholic acid and ursodeoxycholic acid on carrageenan-induced colitis in guinea-pigs. Journal of Pharmacy and Pharmacology 53, 1711–1720 (2001).

78. Wang, Z., Chen, J., Chen, Z., Xie, L. & Wang, W. Clinical effects of ursodeoxycholic acid on patients with ulcerative colitis may improve via the regulation of IL-23-IL-17 axis and the changes of the proportion of intestinal microflora. Saudi Journal of Gastroenterology 27, (2021).

79. Sipos, T. Compositions containing salts of bile acid-aminosalicylate conjugates. (1994).

80. Engevik, K. A. et al. A high-throughput protocol for measuring solution pH of bacterial cultures using UV-Vis absorption spectrophotometry. STAR Protocols 4, 102540 (2023).

81. Scheubert, K. et al. Significance estimation for large scale metabolomics annotations by spectral matching. Nature Communications 8, 1494 (2017).

82. Krassowski, M. ComplexUpset: Create Complex UpSet Plots Using ‘ggplot2’ Components. 1.3.3 10.32614/CRAN.package.ComplexUpset (2020).

83. Gu, Z. circlize: Circular Visualization. 0.4.16 10.32614/CRAN.package.circlize (2013).

84. Dührkop, K. et al. SIRIUS 4: a rapid tool for turning tandem mass spectra into metabolite structure information. Nature Methods 16, 299–302 (2019).

85. Liao, W.-W. et al. A draft human pangenome reference. Nature 617, 312–324 (2023).

86. Zakeri, M., Brown, N. K., Ahmed, O. Y., Gagie, T. & Langmead, B. Movi: A fast and cache-efficient full-text pangenome index. iScience 27, (2024).

87. Gonzalez, A. et al. Qiita: rapid, web-enabled microbiome meta-analysis. Nature Methods 15, 796–798 (2018).

88. Zhu Qiyun et al. Phylogeny-Aware Analysis of Metagenome Community Ecology Based on Matched Reference Genomes while Bypassing Taxonomy. mSystems 7, e00167–22 (2022).

89. Langmead, B. & Salzberg, S. L. Fast gapped-read alignment with Bowtie 2. Nat Methods 9, 357–359 (2012).

90. Zhu, Q. et al. Phylogenomics of 10,575 genomes reveals evolutionary proximity between domains Bacteria and Archaea. Nature Communications 10, 5477 (2019).

91. McDonald, D. et al. Greengenes2 unifies microbial data in a single reference tree. Nature Biotechnology 42, 715–718 (2024).

92. Weng, Y. et al. Calculating fast differential genome coverages among metagenomic sources using micov. Communications Biology 8, 1624 (2025).

