## Supplementary information for "Pan-repository analysis reveals a drug-activating function of microbial bile acid conjugation"

[13]Texas Children's Research Institute, Texas Children's Hospital, Houston, TX, 77030, USA
[14]Helix Chromatography, Prospect Heights, IL, USA
[15]Department of Pathology, Microbiology, and Immunology, Vanderbilt University Medical Center, Nashville, Tennessee, USA
[16]Division of Gastroenterology, University of California, San Diego, La Jolla, CA 92093 USA
[17]Shu Chien-Gene Lay Department of Bioengineering, University of California San Diego, La Jolla, CA 92093 USA
[18]Division of Gastroenterology, Jennifer Moreno Department of Veterans Affairs Medical Center, La Jolla, CA 92093 USA
[19]Chiba University-UC San Diego Center for Mucosal Immunology, Allergy, and Vaccines (CU-UCSD cMAV), La Jolla, CA, 92093, USA
[20]Human Microbiome Research Institute, Cedars-Sinai Medical Center, Los Angeles, CA 90048, USA
[21]Bioinformatics and Systems Biology Program, University of California San Diego, La Jolla, CA, USA
[22]Medical Scientist Training Program, University of California, San Diego, La Jolla, CA, USA
[23]Department of Computer Science and Engineering, La Jolla, CA, USA
[24]Halicioğlu Data Science Institute University of California San Diego, La Jolla, CA, USA
[25]Department of Paediatric Gastroenterology, Royal Hospital for Children and Young People, Edinburgh, UK
[26]School of Medicine, University of Dundee, Dundee, United Kingdom
[27]Strathclyde Institute of Pharmacy and Biomedical Science, University of Strathclyde, Glasgow, G4 0RE, UK
[28]Department of Bioengineering, University of California, San Diego, La Jolla, CA, 92093-0412, USA
[29]Program in Materials Science and Engineering, University of California San Diego, La Jolla, San Diego, CA, 92093, USA
[30]Department of Pediatrics, Vanderbilt University Medical Center, Nashville, TN, USA
[31]Section of Gastroenterology, Hepatology, and Nutrition, Vanderbilt University, Nashville, TN, USA
[32]Division of Pediatric Gastroenterology, Texas Children’s Hospital, Baylor College of Medicine, Houston, TX, USA,
[33]USDA, ARS, Children’s Nutrition and Research Center, Houston, TX, USA
[34]Department of Pharmacology & Immunology, Medical University of South Carolina, Charleston, SC, 29425, USA
[35]Department of Pharmacy Practice & Translational Research, University of Houston, Houston, TX 77004, USA
[36]Wolfe Family Endowed Chair in Microbiome Research at Rady Children’s
[37]Senior Visiting Fellow, Hong Kong University of Science and Technology Jockey Club Institute for Advanced Study

[38]Senior Department of Pathology, University of California, San Diego, La Jolla, California, USA
[39]Hong Kong University of Science and Technology Jockey Club Institute for Advanced Study, Hong Kong University of Science and Technology, Hong Kong SAR, China

*Overall project, data science of the bile acids and discovery.
**For the iPENS and CD-TREAT study:
***For the quantitation data:
****For the mouse interventional study:
*****For the rheumatoid arthritis study:

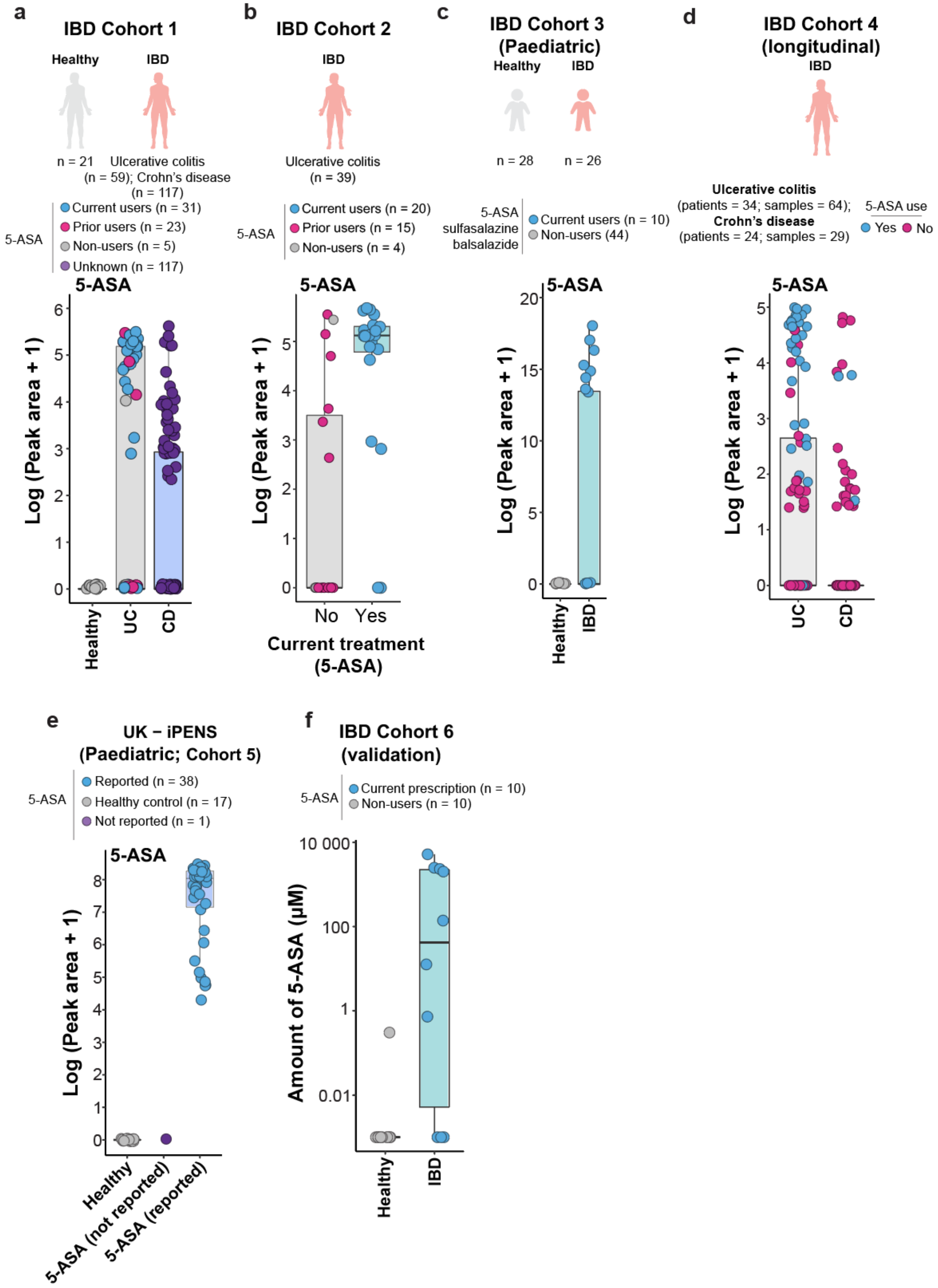
a
IBD Cohort 1
Healthy
IBD
n = 21
Ulcerative colitis (n = 59); Crohn's disease (n = 117)
5-ASA
Current users (n = 31)
Prior users (n = 23)
Non-users (n = 5)
Unknown (n = 117)
5-ASA
Log (Peak area + 1)
Healthy
UC
CD
b
IBD Cohort 2
IBD
Ulcerative colitis (n = 39)
5-ASA
Current users (n = 20)
Prior users (n = 15)
Non-users (n = 4)
5-ASA
Log (Peak area + 1)
No
Yes
Current treatment (5-ASA)
c
IBD Cohort 3 (Paediatric)
Healthy
IBD
n = 28
n = 26
5-ASA sulfasalazine balsalazide
Current users (n = 10)
Non-users (44)
5-ASA
Log (Peak area + 1)
Healthy
IBD
d
IBD Cohort 4 (longitudinal)
IBD
Ulcerative colitis (patients = 34; samples = 64);
Crohn's disease (patients = 24; samples = 29)
5-ASA use
Yes
No
5-ASA
Log (Peak area + 1)
UC
CD
e
UK – iPENS (Paediatric; Cohort 5)
5-ASA
Reported (n = 38)
Healthy control (n = 17)
Not reported (n = 1)
5-ASA
Log (Peak area + 1)
Healthy
5-ASA (not reported)
5-ASA (reported)
f
IBD Cohort 6 (validation)
5-ASA
Current prescription (n = 10)
Non-users (n = 10)
Amount of 5-ASA (μM)
10 000
100
1
0.01
Healthy
IBD

**Supplementary Figure S1 | Detection of 5-ASA across cohorts. a,** Extracted peak area of 5-ASA in healthy volunteers (n = 21), samples from people with ulcerative colitis (n = 59), and Crohn's disease (n = 117). 5-ASA exposure types were color-coded; current users (blue), prior users (pink), non-users (gray), and unknown (purple). **b,** Extracted peak area of 5-ASA in current treatment of 5-ASA of samples from 39 ulcerative colitis patients. 5-ASA exposure types were color-coded; current users (blue), prior users (pink), non-users (gray). **c,** Extracted peak area of 5-ASA in healthy (n = 28) and IBD (n = 26) of a pediatric cohort of IBD. 5-ASA exposure types were color-coded; current users (blue) and non-users (gray). **d,** Extracted peak area of 5-ASA from a longitudinal study of individuals with ulcerative colitis (n = 34; samples 64) and Crohn's disease (n = 24; samples 29). Points are color-coded by 5-ASA exposure; documented 5-ASA use (blue) and not documented use (pink). **e**, Detection of 5-ASA in the untargeted data from the UK iPENS study. The n represents samples. **f**, Concentration (μM) of 5-ASA in 10 individuals without 5-ASA prescriptions and 10 individuals undergoing 5-ASA treatment.

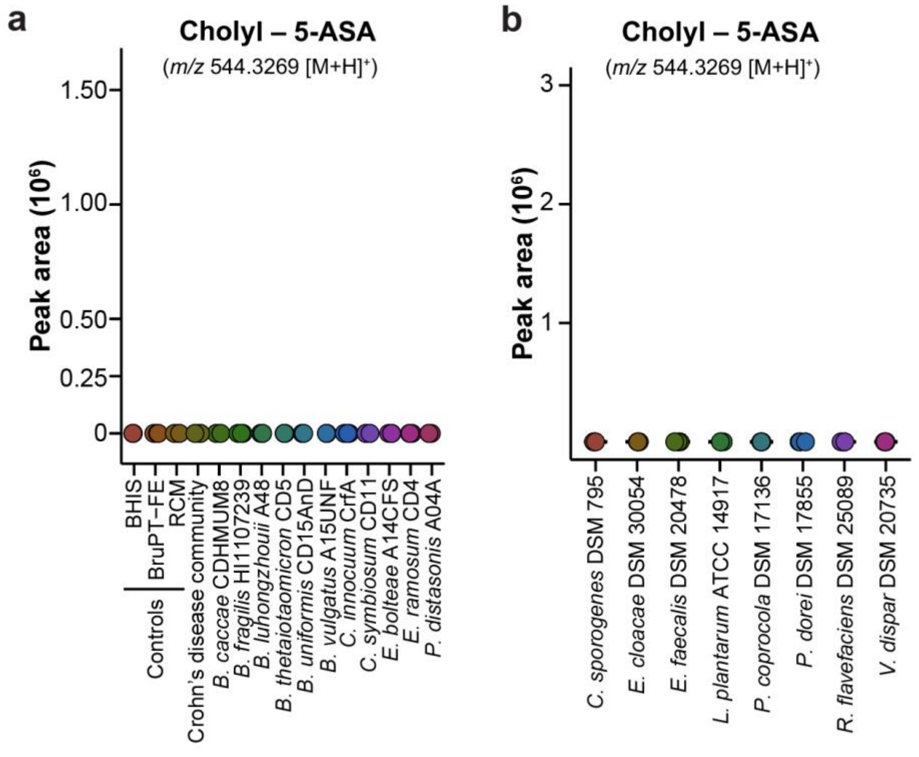


**Supplementary Figure S2 | Human gut microbiota cultures with cholic acid and 5-ASA, and *ex vivo* culture from healthy donors, used for quantitation. a,** Peak area of cholyl-5-ASA measure in monocultures of Crohn's disease-associated isolate incubated with cholic acid and 5-ASA as substrate at baseline (0 h). **b,** Peak area of cholyl-5-ASA measure in monocultures derived from healthy individuals under the same condition than in a.

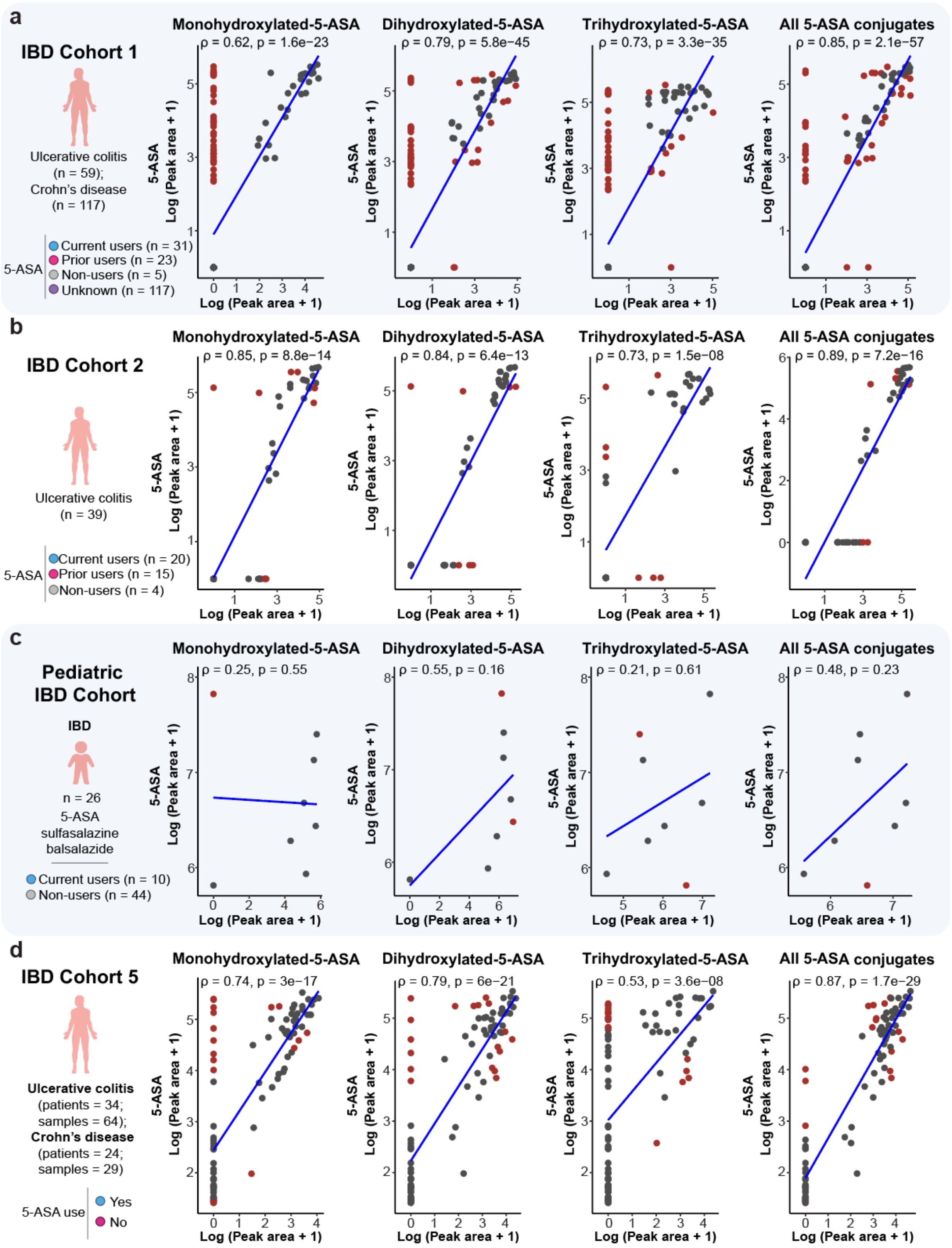
a
IBD Cohort 1
Ulcerative colitis (n = 59); Crohn's disease (n = 117)
5-ASA
Current users (n = 31)
Prior users (n = 23)
Non-users (n = 5)
Unknown (n = 117)
Monohydroxylated-5-ASA
ρ = 0.62, p = 1.6e−23
Dihydroxylated-5-ASA
ρ = 0.79, p = 5.8e−45
Trihydroxylated-5-ASA
ρ = 0.73, p = 3.3e−35
All 5-ASA conjugates
ρ = 0.85, p = 2.1e−57
5-ASA Log (Peak area + 1)
Log (Peak area + 1)
b
IBD Cohort 2
Ulcerative colitis (n = 39)
5-ASA
Current users (n = 20)
Prior users (n = 15)
Non-users (n = 4)
Monohydroxylated-5-ASA
ρ = 0.85, p = 8.8e−14
Dihydroxylated-5-ASA
ρ = 0.84, p = 6.4e−13
Trihydroxylated-5-ASA
ρ = 0.73, p = 1.5e−08
All 5-ASA conjugates
ρ = 0.89, p = 7.2e−16
c
Pediatric IBD Cohort
IBD
n = 26
5-ASA sulfasalazine balsalazide
Current users (n = 10)
Non-users (n = 44)
Monohydroxylated-5-ASA
ρ = 0.25, p = 0.55
Dihydroxylated-5-ASA
ρ = 0.55, p = 0.16
Trihydroxylated-5-ASA
ρ = 0.21, p = 0.61
All 5-ASA conjugates
ρ = 0.48, p = 0.23
d
IBD Cohort 5
Ulcerative colitis (patients = 34; samples = 64);
Crohn's disease (patients = 24; samples = 29)
5-ASA use
Yes
No
Monohydroxylated-5-ASA
ρ = 0.74, p = 3e−17
Dihydroxylated-5-ASA
ρ = 0.79, p = 6e−21
Trihydroxylated-5-ASA
ρ = 0.53, p = 3.6e−08
All 5-ASA conjugates
ρ = 0.87, p = 1.7e−29

**Supplementary Figure S3 | Correlation between 5-ASA and bile acid–5-ASA conjugates abundance across independent IBD cohorts. a-d,** Scatter plots showing the relationship between 5-ASA and monohydroxylated, dihydroxylated, and trihydroxylated bile acid–5-ASA conjugates, as well as the summed abundance of all bile acid–5-ASA conjugates across four independent IBD cohort; **a,** an adult IBD cohort (MSV000084775); **b,** an ulcerative colitis-only cohort (MSV000082094); **c,** a paediatric IBD cohort (MSV000097610); and **d,** a mixed longitudinal IBD cohort (MSV000082221). For each bile acid conjugate class, features corresponding to related isomers were summed prior to analysis. Spearman correlation coefficients (ρ) and two-sided *p* values are shown. Blue lines indicate the linear regression and the red points denote the 20% discordance based on the rank difference between 5-ASA and bile acid–5-ASA conjugates.

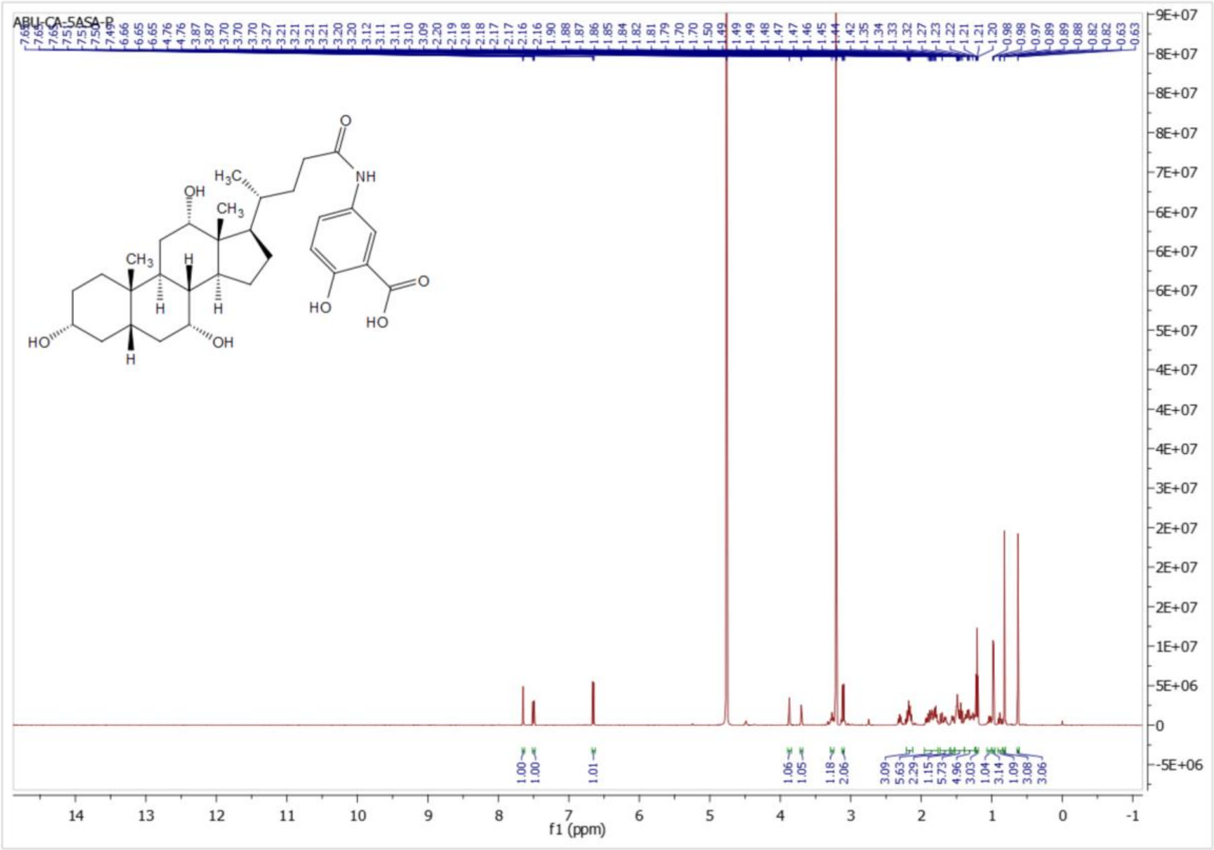


**Supplementary Figure S4 | NMR spectrum of cholyl–5-ASA.**
